# Environmental sensing capacity predicts bacterial ecological strategies and environmental preferences

**DOI:** 10.64898/2026.09.01.748587

**Authors:** Roman S. Altxu, Gauri Shankar, Anna Comas-Pujol, Andrés Francesch-Vázquez, Alberto Pascual-García, Tino Krell, Josep Ramoneda

## Abstract

The ability to sense environmental variation is a prerequisite for ecological success. Sensor domains enable bacteria to detect nutrients, neighboring organisms, and physicochemical conditions, but whether variation in these sensing systems reflects ecological specialization remains unresolved. Here, we analyzed sensor domains across 51,343 bacterial genomes and 255 soil metagenomes spanning a climatic gradient to determine whether sensory repertoires encode bacterial ecological strategies and environmental preferences. Sensory repertoires exhibited strong phylogenetic conservatism and revealed signatures of genome streamlining, indicating that environmental sensing reflects trade-offs associated with maintaining sensory complexity. Taxa occupying environmentally heterogeneous habitats, particularly free-living aerobic generalists, encoded the largest sensory repertoires, consistent with selection for expanded environmental information processing. To link sensory function with ecological adaptation, we mapped experimentally-characterized ligand-binding motifs (LBMs) across genomes and metagenomes. Distinct LBM profiles discriminated host-associated and free-living taxa, aerobic and anaerobic lineages, and generalists and non-generalists, revealing a tight coupling between sensory capacity and ecological strategy. Across soil communities, motifs associated with osmoprotection and oxygen sensing were consistently enriched under increasing aridity, linking sensory function to environmental filtering in natural ecosystems. These findings identify environmental sensing as an important organizational axis of bacterial trait-based ecology that integrates evolutionary history, ecological lifestyle, and adaptation to local conditions. Environmental sensing should be considered for predicting microbial niches and responses to environmental change.

## Introduction

The capacity to perceive and respond to environmental changes is a major determinant of microbial adaptation. Microbial cells continually encounter changes in nutrient availability, redox state, oxygen levels, osmolarity, pH, host signals, and other physicochemical properties of their surroundings [1], making environmental perception fundamental to microbial survival. In order to sense changes in the environment bacteria have evolved a range of different receptor families including sensor histidine kinases, chemoreceptors, adenylate and diguanylate cyclases, cAMP and c-di-GMP phosphodiesterases, and different Ser/Thr protein kinases and phosphatases [2]. Although different in mechanism, these receptors share a similar topology that consists in an input module (or sensor domain) for signal sensing and an output module that generates the corresponding response [3, 4]. These receptors modulate central physiological processes such as chemotaxis, metabolism, or the homeostasis of secondary messenger levels [5] and thus determine the adaptive responses of microbial taxa to environmental contexts.

The capacities for environmental sensing vary drastically across taxa and lead to the expectation that these should define bacterial ecological strategies and habitat preferences [6]. For example, elevated sensory domain diversity has been associated with generalist bacterial lifestyles [7, 8], bacterial virulence [9], and ecological versatility to cope with diverse environmental challenges [10]. The lack of sensory systems has in contrast been associated with oligotrophic lifestyles often occurring in taxa that underwent the process of genome streamlining [11]. Sensor domain repertoires can also be used as ‘fingerprints’ that distinguish microbial communities inhabiting contrasting ecosystems such as different host-associated habitats and discriminate among disease states from gut microbiomes [12]. However, while we keep improving our understanding of the strategies and genes for bacterial stress tolerance and metabolism [13], we lag behind in the identification of the environmental sensing capabilities of taxa, which are likely as important in determining bacterial environmental preferences and thus for predicting responses to environmental change.

Bacterial environmental sensing relies on signal transduction systems that convert environmental cues into regulatory responses affecting metabolism, motility, stress tolerance, and other cellular processes [4]. Among these, two-component systems are widespread signalling architectures, typically comprising a sensor histidine kinase and a cognate response regulator connected through reversible phosphorylation [14, 15]. Histidine kinases typically detect chemically diverse molecules and physical stimuli through sensory domains that show large variation in topology and molecular mechanism [16]. Whereas most two-component systems modulate gene expression in response to signals, other families modulate cAMP and c-di-GMP second messenger levels, bind to RNA or control protein phosphorylation [17]. Extracellular or periplasmic sensor domains can directly recognize environmental cues like small molecules, whereas cytosolic sensory domains detect intracellular as well as extracellular signals [14]. However, the functionality of 98.7% of histidine kinase sensor domains remains unknown [12], and sensor domain composition alone provides limited information about the environmental signals that individual receptors can detect. A large body of research is dedicated to the description of sensor protein structures and their ligand-binding functionality [6, 18–21], such that it might be feasible to establish testable linkages between bacterial sensory capacity, ecological strategies, and environmental preferences. However, due to their fast evolutionary pace [22], the signal recognised by sensor domains is not reflected in overall sensor domain sequence similarity [23], which in turn hampers the functional annotation of uncharacterised receptors by sequence similarity with characterised receptors.

Experimentally-characterized sensor domains provide an initial basis for establishing the connection between the sensory capacity of bacteria and environmental preference. For example, FAD-binding domains from the PAS superfamily, such as that of the *E. coli* Aer receptor, can function as intracellular redox-sensing modules that couple changes in cellular respiration to behavioural responses such as aerotaxis [24, 25]. Among extracytosolic sensory systems, Cache domains constitute a major superfamily of ligand-binding sensor domains, with characterized members recognizing chemically diverse compounds such as amino acids [26], biogenic amines [18, 27], sugars [28], inorganic ions [29], organic acids [30], purine derivatives [19], quorum-sensing signals [31], and other ligands [32–34]. These experimentally established ligand-recognition functions provide a mechanistic basis for interpreting the ecological distribution of sensor domains. For example, biogenic amines such as choline and betaine can function as compatible solutes under osmotic stress [35], as do some amino acids [36]. Besides choline, the transcriptional regulator BetI has been shown to sense the compatible solutes trimethylamine and acetylcholine, which also had osmoprotective functions [37]. This leads to the expectation that their recognition should be adaptive in environments with elevated solute concentrations, such as in saline systems or dry soils. Such functions may become particularly relevant under environmental gradients associated with global change, such as aridity gradients, where declining water availability is accompanied by increased osmotic stress [38, 39]. Experimentally characterized ligand-binding functions thus provide a means to generate mechanistically informed predictions about the sensory capabilities of taxa, their environmental associations, and how they might respond to environmental change.

The overarching goal of this study was to determine the role of environmental sensing in the ecological adaptation of bacteria across habitats and environmental conditions. Because environmental sensing can be examined at both the levels of individual genomes and microbial communities, we combined information at the single-species level, enabling exploration of phylogenetic context and taxonomic relationships (51,343 bacterial genomes), with community-level information, which enables associations with broader environmental drivers (255 soil metagenomes across a broad aridity gradient). To this end, we compiled a database of 84,881 histidine kinase sensor domain variants and 13 ligand-binding motifs which allowed us to 1. determine the taxonomic and phylogenetic distribution of bacterial sensor domain diversity, prevalence, and ligand-binding functionality across the main bacterial phyla; 2. establish associations between sensor domain diversity and ligand-binding functionality with bacterial lifestyles, ecological strategies, and environmental preferences; and 3. showcase the interest and utility of investigating bacterial environmental sensing capabilities for understanding microbial community responses to environmental change. While environmental sensing has been largely excluded from trait-based analyses of microbial communities due to the limited knowledge on sensor domain variation and functionality, we show that sensor domains and their ligand-binding functions represent an important trait axis that can be leveraged to expand our trait-based understanding of bacterial environmental adaptation.

## Materials and Methods

### Data acquisition

In this study we utilized a protein database covering the known bacterial histidine kinase sensor domain sequence space [12]. This database is expected to be generally representative of the known sensor domain variability in bacteria as it was obtained from a global compilation of >20,000 metagenomes covering 76 different biomes [12], and histidine kinases are the most common form of two-component sensory systems in bacteria. For all 113,186 sensor domains contained in the original database, we first filtered out those predicted to be intracellular sensor domains using the TMHMM v2.0 protein topology prediction tool [40], as we were interested in investigating those domains involved in sensing of the extracellular environment. We however retained intracytosolic PAS domains for being widely distributed and for having well-described sensory functions [41], including the sensing of oxygen and the cellular redox potential that reflects in turn the metabolic state and nutrient availability in the environment. For the inferred extracytosolic domains, we classified them into protein families with HMMER v3.1b2 [42], using the hmmscan command on the Pfam database v38.0 [43]. In this way, we obtained a database of bacterial extracytosolic sensor domains and could subsequently investigate it across genomes and environmental metagenomes.

To explore the taxonomic distribution of bacterial sensor domains, we compiled genomic data from ∼143,000 unique bacterial taxa (“species clusters”) available in the Genome Taxonomy Database (GTDB) (release 226; [44]). We then carried out a quality filtering step to include only genomes with estimated >90% completeness and <5% contamination based on CheckM [45]. We also kept only those containing an assembled 16S rRNA gene, yielding a total of 51,343 high-quality genomes (Supplementary Dataset 1). To explore the distribution of bacterial sensor domains across an environmental gradient, we retrieved publicly available soil metagenomic data from the Australian Microbiome Initiative Biomes of Australian Soil Environments (BASE) project [46], from which we selected 255 soil shotgun metagenomic samples representing a broad range of locations and climatic conditions across the Australian continent (mean annual precipitation 138-2959 mm). We selected this dataset to cover the broadest possible range in aridity, as it drives water availability which is a key determinant of soil bacterial survival [47]. These samples were all obtained from 0-10 cm depth from natural areas (designated “natural and conservation areas” in BASE). We downloaded the associated soil property metadata including conductivity, pH, and environment category, which we complemented by assigning the GPS coordinates to the aridity index from the Global Aridity Database version 3 map at 30 arc-second resolution [48]. The sample identifiers and metadata are detailed in Supplementary Dataset 2, and the raw sequencing data are available at the Australian Microbiome Initiative Data Portal (https://data.bioplatforms.com/organization/about/australian-microbiome).

### Metagenome assembly and taxonomic profiling

We retrieved raw reads from the BASE project database and performed quality filtering by removing adapters and other putative contaminant sequences using trimmomatic v0.39 [48] (parameters: ILLUMINACLIP:TruSeq3-PE.fa:2:30:10, LEADING:3, TRAILING:3, SLIDINGWINDOW:4:15, MINLEN:36), and checking their quality using FASTQC [49]. We assembled the processed reads into metagenomes by running MEGAHIT v1.2.9 [50] on the ‘meta-large’ preset, and assessed the quality of the assembly using QUAST v5.0.2 [51]. The protein-coding regions of the assembled metagenomes were later predicted using Prodigal v2.6.3 [51] on ‘-p anon’ mode. We performed taxonomic profiling of the communities using SingleM v0.20.3 [52] (Metapackage database S5.4.0) ‘pipe’ command on forward and reverse raw metagenomic reads, and processed the obtained mOTU tables using SingleM ‘summarise’ option which allowed us to extract relative abundances of taxa across different taxonomic levels.

### Bacterial sensor domain annotation

The annotation and profiling of sensor domains in a given genome or metagenome was carried out through mapping of their translated predicted protein-coding region (Prodigal v2.6.3; [51]) to our filtered sensor domain database. To this end, we ran DIAMOND v2.1.14 [52] using blastp option, with ‘–-very-sensitive’ preset and reporting only the best match below an e-value cutoff of 0.001. In metagenome assemblies, we took all read mapping putative genes matching a given sensor domain, selecting the best match for every given read. To avoid low-quality matches, we only considered predicted gene-sensor domain matches with >50% subject sequence cover and bitscore >50. We also filtered out rare domains appearing in less than 10 samples across the metagenomic dataset and a mean read counts per million (CPM) below 1. For high-quality genomes retrieved from GTDB, we only considered protein matches with a subject cover value >90% and bitscore >50. The abundance of every given sensor domain in each genome was calculated as the sensor domain gene count normalized by the total number of protein entries in the genome.

### Identification of ligand binding motifs within sensor domains

To elucidate the putative ligands of our bacterial sensor domain collection, we carried out a literature search for experimentally described ligand binding motifs across the main histidine kinase sensor domain families [21]. The families with available ligand binding motifs were GAF (1 motif, [53]), PAS (9 motifs, [53]), and dCache (3 motifs, [26]; [18]; [32]. From the collected ligand binding motifs, we constructed regular expression patterns for each family and searched for them in our filtered sensor domain database, which allowed us to identify their putative ligands. The full database of sensor domain families with their described ligand binding motifs can be found in Supplementary Table S1.

### Ecological and phylogenetic analysis

We obtained annotations for bacterial habitats, lifestyles, and environmental preferences (oxygen, temperature, pH, and salinity) from metaTraits [54, 55]. This database contains bacterial trait annotations for bacterial strains and MAGs whose identity can be directly linked to GTDB accessions. In order to analyze the phylogenetic distribution of sensor domains across bacterial taxa, we obtained a robust phylogeny of our genomes by trimming the tree from GTDB which is based on the alignment of 120 single-copy near-universal marker genes [44]. We tested whether differences in the presence/absence of sensor domains between genomes based on the Jaccard dissimilarity index correlated with cophenetic distances between them using Mantel test (9999 permutations, *vegan* R package v2.7-3; [56]). We then estimated phylogenetic signal in the composition of ligand binding motifs by coding the presence/absence of motifs for each individual ligand as a binary trait and then using the phylogenetic *D* index for binary traits [57], implemented in the *ape* R package (v5.7-1; [58]). We visualized and edited the tree using *iTOL* (v5; [59]) by selecting a single random representative genome from each family within the 11 most abundant bacterial phyla.

### Statistical analysis

We used linear regression to test the association between bacterial genome size and the prevalence of sensor domains within genomes. Shapiro-Wilk tests indicated the data on the prevalence of both sensor domains and ligand binding motifs in them were overall non-normally distributed. We accordingly tested for pairwise differences between groups using Mann Whitney U tests throughout the manuscript, and tested for the influence of aridity on the composition of ligand binding domains in metagenomes using Kruskal-Wallis tests. We tested for differences in the prevalence of LBMs across bacterial ecological categories using linear regression, and represented these differences using z-scores to avoid sampling biases. We used Spearman’s rank correlations to establish relationships between the relative abundance of particular bacterial phyla and the prevalence of individual sensor domains across metagenomes. We corrected P-values for multiple comparisons using Benjamini-Hochberg correction. Phyla were assigned an ‘aridity preference’ by dividing the difference of mean relative abundance under arid and humid conditions by the mean relative abundance under humid conditions.

We represented the sequence similarity space of sensor domains embedding our sequences using ESM-2 Transformer protein language model (‘esm2_t12_35M_UR50D’; [60]), and projecting the sequence embedding using t-SNE at perplexity values of 0.5%, 1%, 5% and 10% of our dataset size to capture local and global similarities of our sequence space. To test whether the capacities for binding particular ligands co-occurred more frequently than expected across bacterial genomes, we used Fisher’s exact tests on all pairwise comparisons of ligand binding motifs. In order to test the dominant environmental factors underlying differences in the sensor domain composition of metagenomes between arid (AI = 0-0.65) and humid (AI > 0.65) sites, we used distance-based redundancy analysis (db-RDA) on Bray-Curtis distances using *vegan* R package (v2.7-3; [56]). We used a Support Vector Machine (SVM) classifier to determine the sensor domains that best distinguished soil metagenomes from arid and humid sites. We used hierarchical clustering to represent differences in the sensor domain profiles of metagenomes, and represented these differences using Principal Components Analysis (PCA). All statistical analyses were performed in base R (v4.5; R Core Team, 2025) and otherwise as specified.

## Results and Discussion

### A metagenomic approach to investigate bacterial environmental sensing capacity across taxa and environmental gradients

Although the sequence space of bacterial sensor domains is relatively well characterized, the taxonomic and environmental distributions of most of these proteins remain unknown. We used the histidine kinase sensor domain database described by [12], which captures the known sequence space of the most widespread bacterial environmental sensing strategy. We focused on extracytosolic domains involved in sensing primarily extracellular signals and thus directly implicated in environmental sensing [21], while also including intracellular PAS domains because of their well-established sensing functions [41]. Accordingly, we filtered the reference database from 113,186 to 84,881 sequences corresponding to extracytosolic and PAS domains (Fig. 1).

**Figure 1.**
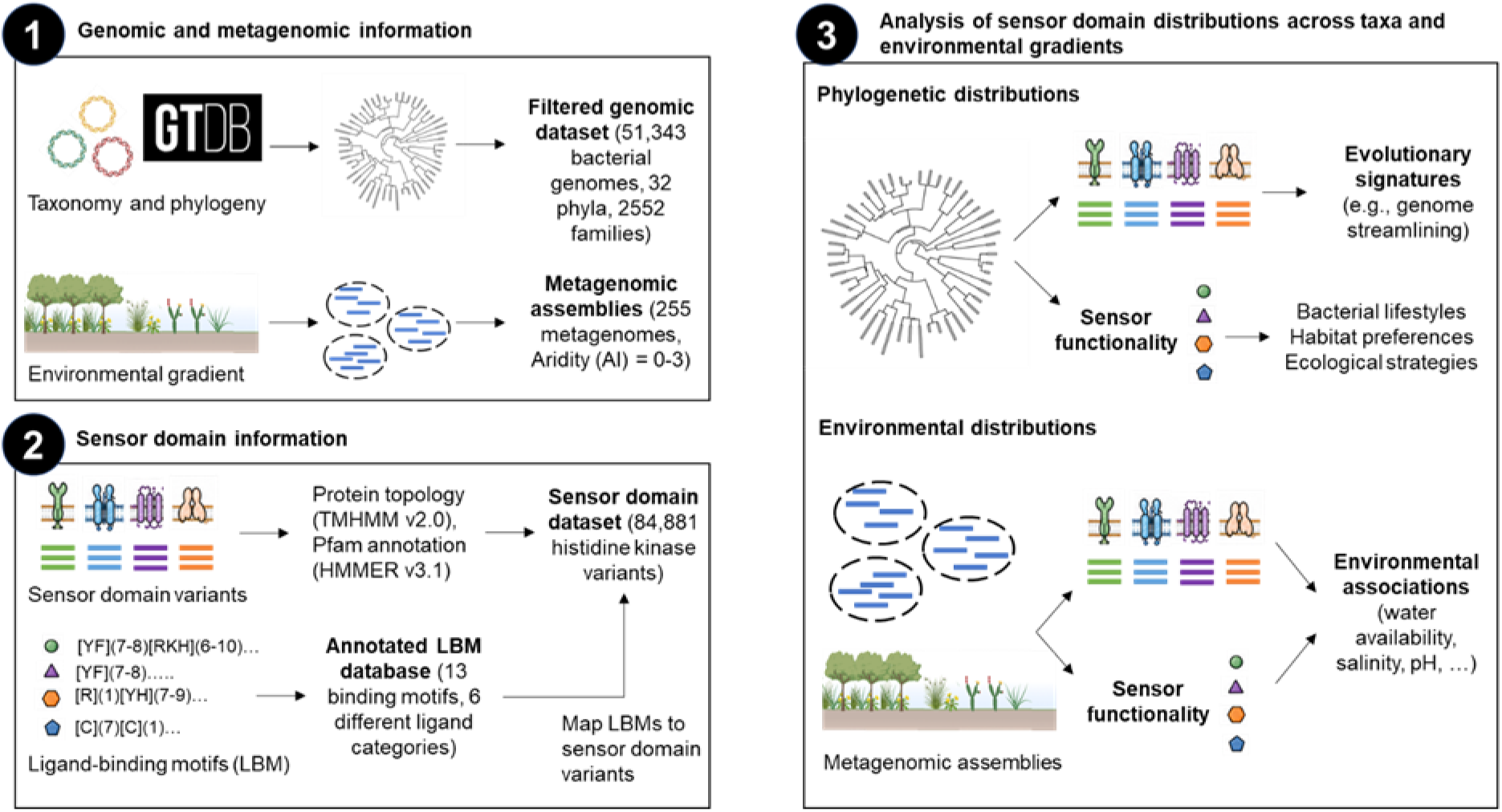
A bioinformatics approach to investigate the taxonomic and environmental distribution of sensor domains in bacteria. We filtered a database of 113,186 bacterial sensor domain variants to obtain only extracytosolic domains. We then mapped those variants to bacterial genomes as well as to metagenomic contigs from metagenomes of soil microbial communities across a broad climatic gradient. We then screened the literature for sensor proteins whose binding ligands had been experimentally linked to their ligand binding motifs (LBMs), as the ligand-binding capacities of sensors underlie their functional roles in environmental sensing. We then screened our mappings for these LBMs and evaluated their distribution across bacterial taxa and phylogenetic groups, bacterial habitats, lifestyles, and environmental preferences, and soil metagenomes across a model aridity gradient.

Because sensor domain functions are largely determined by their binding ligands, we compiled information on described ligand-binding motifs (LBMs) for 5 protein families and 284 subfamilies represented in our database. We focused on the highly represented GAF, PAS, and dCache families, excluding the abundant CHASE domains because no characterized LBMs could be identified (Supplementary Table S1). We also identified 26 putative extracytosolic PAS domains that likely belong to the sCache family [27]. Our search identified 13 LBMs associated with FAD, heme groups, 4-hydroxycinnamic acid, amino acids and biogenic amines, purines, divalent cations, and carboxylates (Supplementary Table S1). Although knowledge of bacterial sensor domain LBMs remains limited to a small number of well-characterized representatives, the ligands represented here span diverse functions with potentially testable links to bacterial lifestyles and environmental preferences (Supplementary Table S1; Fig. 1).

### Taxonomic distribution of bacterial sensor domains

To investigate the distribution of histidine kinase sensor domains across bacteria (hereafter called ‘sensor domains’ for brevity), we obtained 51,343 high-quality genomes from GTDB spanning 32 phyla and 2,552 families (Supplementary Data 1). Our genomic dataset contained 25,147 unique sensor domain variants, representing 29.6% of the reference database. Given the rapid evolution of bacterial sensor proteins [21, 22], we first examined the lineage specificity of sensor repertoires. Analysis of the most represented phyla showed that 71.9% of domains were phylum-specific (Fig. 2A). Consistently, sensor domain composition was positively associated with phylogenetic distance (Mantel test, r = 0.30, P = 0.0001; Supplementary Fig. S1). Among domains present within a given clade, the prevalence of unique domains decreased from 27.8% at the phylum level to 12.8% at the class level and 2.9% at the genus level (Supplementary Fig. S2). Thus, closely related bacteria tend to possess more similar sensory repertoires, whereas increasing evolutionary distance is associated with greater differentiation. This lineage-associated organization is consistent with the evolutionary diversification of histidine kinases and two-component systems [61–63], and with the phylogenetically structured distribution of Cache domains across prokaryotes [27].

**Figure 2.**
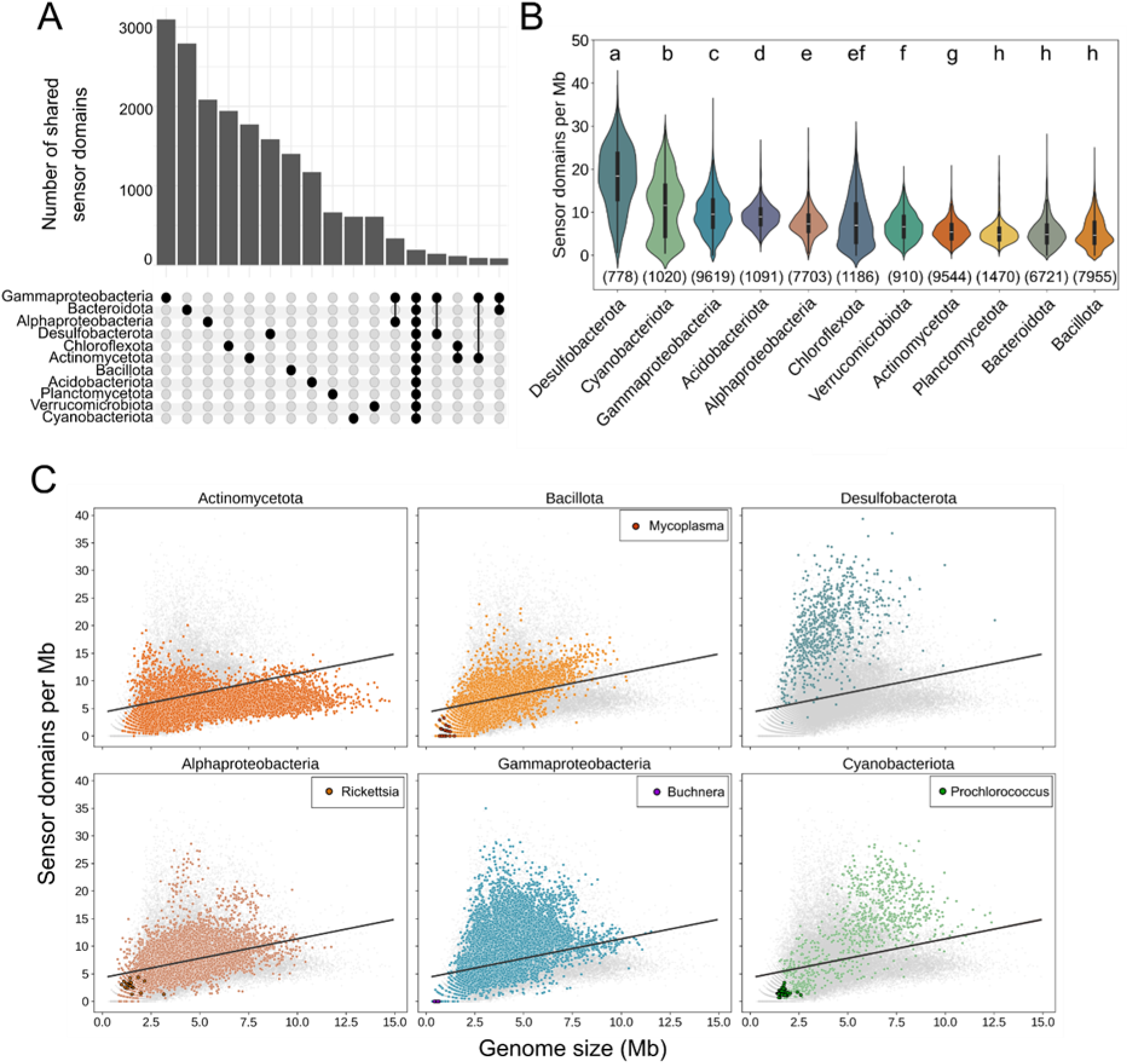
Prevalence of sensor domains across the main bacterial phyla and implications for genome streamlining. (A) Distribution of unique and shared sensory domains across genomes of the main bacterial phyla. (B) Prevalence of sensory domains calculated as number of sensory domains per megabase (Mb) across bacterial phyla. (C) Relationship between the prevalence of sensory domains and genome size. Black lines indicate the linear regression of the relationship. Genomes belonging to well-described groups with streamlined genomes are indicated on the plots: *Rickettsia* and *Mycoplasma* (endoparasites), *Buchnera* (endosymbiont), and *Prochlorococcus* (free-living). N = 49711 genomes.

Despite this phylogenetic structuring, we identified 224 sensor domains shared across all selected phyla (0.89% of the dataset; Fig. 2A). We therefore examined whether these broadly distributed domains also shared sequence features. Sequence-based t-SNE revealed a partially cohesive cluster, with 28.6% of the shared domains grouping together (Supplementary Fig. S3). This suggests that a subset of sensor domains retains detectable sequence similarity across deeply divergent bacterial lineages. Similar coexistence of conserved sequence features and lineage-specific diversification has been reported for bacterial chemoreceptors, including dCache receptors that share related sensory domains while differing in ligand-recognition properties [64]. Indeed, sensor domains can combine conserved structural features with extensive variation in ligand-binding regions [21], such that sequence similarity does not necessarily imply functional similarity. Our dataset provides a foundation for investigating the evolutionary processes underlying these patterns across a broader taxonomic scale than previous studies [8, 65].

### Prevalence of sensor domains reveals signatures of genomic streamlining

The prevalence of sensor domains (number of sensor domains per Mb of genome sequence) varied markedly across bacterial phyla (Fig. 2B). Phyla characterized by broad ecological distributions and metabolic versatility, including the Desulfobacterota [66], Cyanobacteriota [67], class Gammaproteobacteria [68], and Myxococcota [69], encoded the largest sensory repertoires, whereas Planctomycetota, Bacteroidota, and Bacillota displayed a lower prevalence of sensor domains. These observations are consistent with the proposed role of environmental sensing diversity in coping with environmental complexity [7, 10], and are similar to the patterns observed for these phyla in deep subsurface environments [70]. There was however high variation in the prevalence of sensor domains within all phyla (Fig. 2B), likely reflecting variation in lifestyles and habitat preferences within these groups as has been previously reported within the Cyanobacteriota [71]. We verified that isolates and MAGs have a similar prevalence of sensor domains, so the cultivability of taxa is unlikely to be related to the prevalence of sensor domains (Supplementary Fig. S4).

Across all genomes, sensor domain prevalence was positively associated with genome size (Spearman’s ρ = 0.44, P < 0.001; Fig. 2C). This relationship enabled the identification of taxa with larger or smaller sensory repertoires than expected based on genome size, which has been previously identified as a potential signature of genome streamlining [7]. Several phyla, including Desulfobacterota, Cyanobacteriota, and Gammaproteobacteria, were enriched in sensor domains relative to genome size, whereas Actinomycetota were predominantly depleted (Fig. 2C). We next examined bacterial groups with well-established signatures of genome streamlining. Free-living oligotrophs (*Prochlorococcus*) as well as symbiotic and parasitic taxa (*Buchnera*, *Rickettsia*, and *Mycoplasma*) consistently encoded fewer sensor domains than expected from genome size, with some genomes lacking sensor domains altogether (Fig. 2C; Fig. S5). These results indicate that environmental sensing capacity is strongly reduced during genome streamlining [72], whether driven by selection for genomic efficiency in nutrient-poor environments [73] or by reductive evolution associated with stable host-associated lifestyles [74].

### Phylogenetic distribution of the ligand binding capacities of bacterial sensor domains

To better characterize bacterial sensory capacity, we investigated the prevalence of experimentally characterized ligand-binding motifs (LBMs) across bacterial genomes (Fig. 3; Supplementary Fig. S6; Table S1). We identified 156 sensor domains carrying characterized LBMs, distributed across 16,172 genomes. The most represented motifs were associated with 4-hydroxycinnamic acid (p-coumaric acid, a plant defense compound; [75]), biogenic amines (osmoprotectants such as acetylcholine, choline, and betaines; [18]), and carboxylates (Supplementary Fig. S6), although their distribution differed markedly among protein families (Supplementary Fig. S7). We also identified motifs involved in sensing redox and oxygen-related compounds (FAD and heme), divalent cations, amino acids, and purines (Table S1). Among the domains with known LBMs, the most abundant subfamily was that of the amine-sensing dCache domains (85.5% of dCache domains with LBMs; 59 proteins exclusively found within this family), whereas 4-hydroxycinnamic acid- and carboxylate-binding motifs were most prevalent in PAS domains (29.9% and 27.6%, corresponding to 26 and 24 proteins, respectively; Supplementary Fig. S7). Together, these ligands span diverse physiological functions, providing a suitable framework to investigate links between sensing capacity, phylogeny, lifestyles, and environmental preferences.

**Figure 3.**
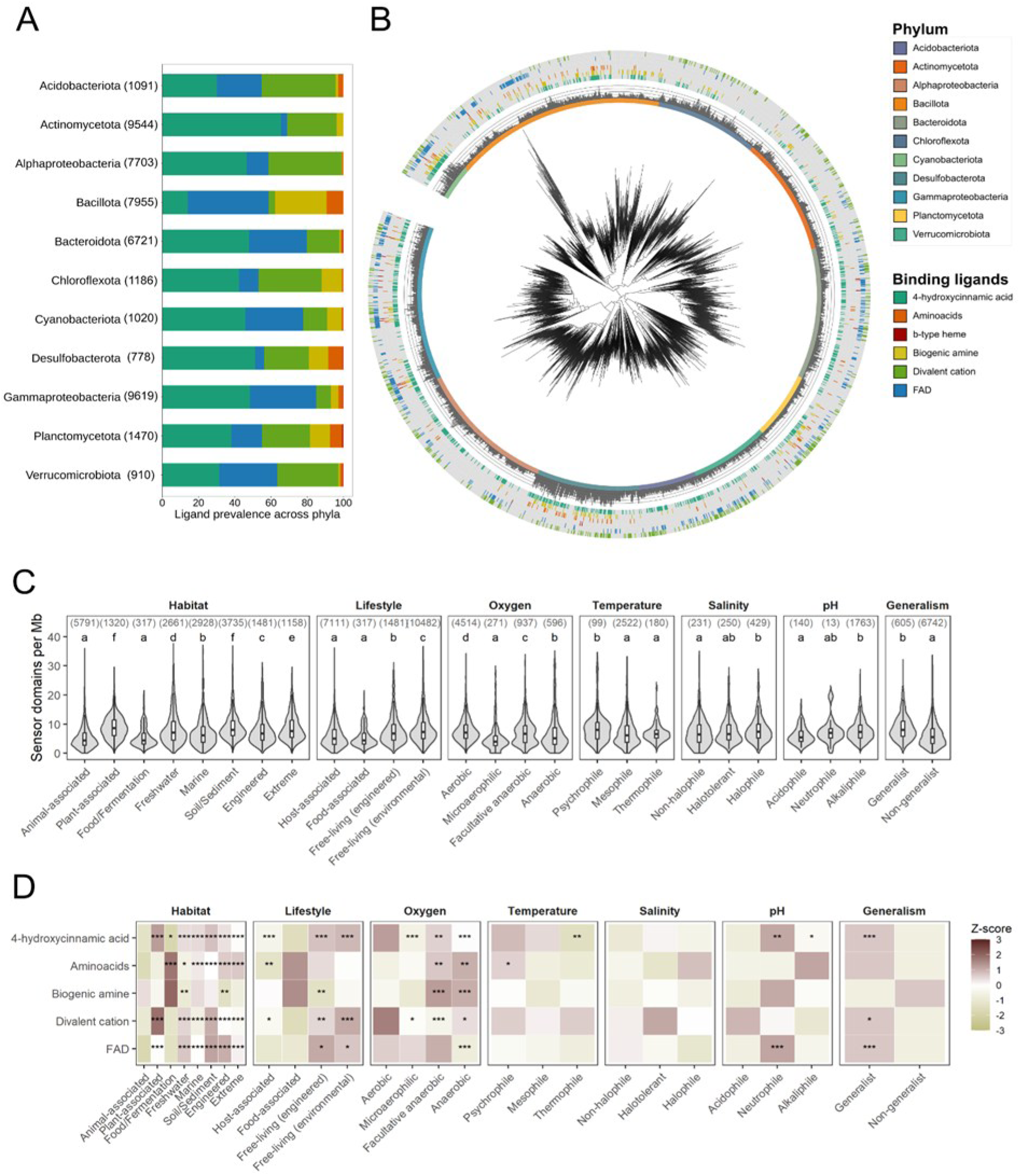
Taxonomic, phylogenetic, and environmental distribution of experimentally characterized ligand binding motifs. (A) Distribution of experimentally characterized ligand binding motifs (LBMs) across bacterial phyla. (B) Phylogenetic tree depicting the prevalence of sensor domains and experimentally characterized ligand binding motifs across the main bacterial phyla. (C) Prevalence of sensor domains calculated as number of sensor domains per megabase (Mb) across bacterial habitats, lifestyles, and environmental preferences. (D) Ecological associations of bacterial LBMs across bacterial habitats, lifestyles, and environmental preferences. For visualization purposes the phylogeny includes a single genome representative from each of the 2190 bacterial families belonging to the 11 most represented phyla in the Genome Taxonomy Database (GTDB). Inner circle: Phylum affiliation; middle circle: sensor domain prevalence expressed as density (number of sensory domains per megabase); outer circles: presence of ligand binding domains across genomes (see legend). The tree was constructed by trimming the GTDB phylogeny (release 226). Letters in panel C indicate statistical differences between groups based on Benjamini-Hochberg corrected P-values from Mann Whitney U tests (significance set at P < 0.05). Number in brackets depict the total number of genomes (N) in each category. Asterisks in panel D depict significant enrichment of LBMs across categories based on generalized linear models with Benjamini-Hochberg P-value correction (*P < 0.05, **P < 0.01, ***P < 0.001).

LBMs were unevenly distributed across bacterial phyla (Fig. 3A). FAD- and b-type heme-binding motifs occurred predominantly in Gammaproteobacteria (45.6% and 51.2% of genomes containing these motifs, respectively), whereas amino acid- and biogenic amine-binding motifs were most common in Bacillota (38.1% and 48.5%). Motifs recognizing 4-hydroxycinnamic acid and divalent cations were most prevalent in Alphaproteobacteria (15.9% and 32.4%) and Actinomycetota (13.0% and 12.3%) (Fig. 3A). Although carboxylate-binding motifs were the third most abundant motif type in the reference database (Supplementary Fig. S6), they were detected in only 75 of 51,343 genomes and were therefore excluded from subsequent genomic analyses. Importantly, isolate genomes and MAGs showed comparable LBM compositions (Supplementary Fig. S8), suggesting limited taxonomic bias associated with culturable organisms.

Despite these differences, most LBMs exhibited broad phylogenetic distributions, and motif composition varied substantially among taxa within the same phylum (Fig. 3B). Divalent cation- and 4-hydroxycinnamic acid-binding domains were particularly widespread, whereas FAD-, amino acid-, biogenic amine-, and heme-binding motifs showed more localized distributions. Consistent with this pattern, only FAD-binding motifs displayed relatively strong phylogenetic conservatism (D = 0.324, P < 0.001), while motifs binding 4-hydroxycinnamic acid, amino acids, and biogenic amines exhibited weaker but significant phylogenetic clustering (Table S2). FAD-binding motifs were especially concentrated within several clades of Bacillota, Bacteroidota, and Gammaproteobacteria (Fig. 3B). Notably, Bacillota displayed a distinct LBM repertoire, characterized by the combined presence of FAD-, amino acid-, and biogenic amine-binding motifs, consistent with their separation in the sequence-level analysis (Supplementary Fig. S3). We also detected significant co-occurrence of FAD- and 4-hydroxycinnamic acid-binding motifs with motifs for other ligands (Table S3), indicating that most taxa are equipped to sense multiple classes of environmental stimuli rather than specializing on individual ligands.

### Associations between sensor domain prevalence and ligand binding capacity with bacterial lifestyles and environmental preferences

We then tested whether environmental sensing capacity, measured as sensor domain prevalence and ligand-binding repertoires, was associated with bacterial lifestyles and environmental preferences. Both metrics varied strongly across habitats, lifestyles, and preferences for oxygen, temperature, pH, and salinity (Fig. 3C,D). Environmental free-living taxa harbored more sensor domains than host-associated taxa (8.33 ± 4.95 vs 5.97 ± 4.07 sensor domains Mb⁻¹; Kruskal-Wallis χ²(3) = 1326.77, P < 0.001; Fig. 3C). Likewise, taxa associated with soils and sediments (9.11 ± 4.60 sensor domains Mb⁻¹), plant environments (8.88 ± 3.85 sensor domains Mb⁻¹), and extreme environments (8.77 ± 4.53 sensor domains Mb⁻¹) showed higher sensor domain prevalence than animal-associated (5.31 ± 3.81 sensor domains Mb⁻¹) and food-associated taxa (5.47 ± 3.22) (Kruskal-Wallis χ²(7) = 2794.78, P < 0.001). These observations confirm the main hypothesis that taxa from more complex or temporally variable environments require a broader sensory repertoire to navigate their surroundings [76]. Host- and food-associated taxa often also lack biosynthesis genes for essential metabolites [77–79], which can be explained by the Black Queen Hypothesis of reductive evolution [80] and can also explain the reduced sensory capacities of host-associated taxa. Together, these observations identify environmental sensing capacity as a genomic trait strongly affected by genome streamlining.

Sensor domain prevalence also varied across environmental preferences. Aerobes (7.75 ± 3.85 sensor domains Mb⁻¹) and facultative anaerobes (7.09 ± 4.32 sensor domains Mb⁻¹) encoded more sensor domains than obligate anaerobes (7.04 ± 6.17 sensor domains Mb⁻¹) (Kruskal-Wallis χ²(3) = 253.52, P < 0.001), while habitat generalists had substantially larger sensory repertoires than non-generalists (8.74 ± 4.16 vs 6.33 ± 4.12 sensor domains Mb⁻¹; Mann-Whitney P < 0.001) (Fig. 3C). Given the strong temporal variability of oxygen concentrations even in typically oxic environments such as surface soils [80], expanded sensory repertoires may facilitate the continuous behavioral and metabolic adjustments required for ecological success [81]. More broadly, the enrichment of sensor domains among habitat generalists suggests that sensory capacity contributes to bacterial ubiquity [55], complementing previously identified genomic predictors such as genome size and transcription-factor diversity [82].

We next investigated whether ligand-binding capacities reflected bacterial lifestyles and environmental preferences. The capacities for binding 4-hydroxycinnamic acid, FAD, and divalent cations clearly distinguished free-living from host-associated taxa, generalists from non-generalists, and aerobic from anaerobic taxa (Fig. 3D). Anaerobes were the only group significantly depleted in FAD-binding motifs (Z-score = −1.44, P < 0.001). Because FAD-containing PAS domains act as major redox and oxygen sensors in bacterial aerotaxis systems [83], these results indicate that ligand-binding capacities capture ecologically relevant physiological differences among taxa. Current efforts to predict oxygen preferences from genomes often rely on amino-acid composition [84], oxygen-utilizing enzymes [85], or protein-family annotations [86]. Our results show that information on ligand-binding motifs alone can also discriminate anaerobic taxa and may help infer ecological traits in the many bacteria that remain poorly characterized.

Habitat-specific comparisons revealed distinct sensory profiles. Plant-associated taxa were enriched in divalent cation-binding motifs (Z-score = 1.73, P < 0.001) and 4-hydroxycinnamic acid-binding motifs (Z-score = 1.18, P < 0.001), whereas food-associated taxa showed the strongest depletion of 4-hydroxycinnamic acid-binding motifs (Z-score = −1.74, P = 0.023). Food-associated taxa were, in turn, enriched in amino-acid-binding motifs (Z-score = 1.65, P < 0.001), contrasting with freshwater taxa, which were depleted in these motifs (Z-score = −0.63, P = 0.013) (Fig. 3D). These patterns mirror previously reported differences in amino-acid biosynthetic capabilities across habitats. Food-associated communities are often enriched in amino-acid auxotrophs, whereas freshwater communities are strongly depleted in such taxa [79]. Thus, the observed distribution of amino acid-binding motifs may reflect differences in the ecological importance and availability of extracellular amino acids across environments [23].

Ligand-binding capacities also differentiated taxa sharing similar lifestyles but occupying different environments. Among free-living bacteria, freshwater taxa were enriched in FAD-binding motifs (Z-score = 0.68, P < 0.001) and divalent-cation-binding motifs (Z-score = 0.49, P < 0.001), whereas both motif types were depleted in marine taxa (Z-score = −0.04, P < 0.001 and Z-score = −0.67, P < 0.001, respectively) (Fig. 3D). In contrast, marine taxa were enriched in amino-acid-binding motifs (Z-score = 0.22, P < 0.001) and showed a positive trend for biogenic-amine-binding motifs (Z-score = 0.46, P = 1.00), whereas freshwater taxa were depleted in both categories (Z-score = −0.63, P = 0.013 and Z-score = −0.85, P = 0.002, respectively). Although these habitat-specific associations should be interpreted cautiously, dissolved amino acids and amines constitute important components of marine dissolved organic matter and can provide major sources of carbon, nitrogen, and osmoprotective compounds for microorganisms [87]. The ability to detect amino acid-rich nutrient patches may therefore confer a competitive advantage to marine bacterioplankton [88, 89]. Conversely, the enrichment of divalent-cation-sensing motifs in freshwater taxa is consistent with the greater spatiotemporal variability of ionic conditions in freshwater systems, subject to local geological and hydrological conditions [90], relative to the relatively stable ionic composition of seawater, subject to mixing with a near-constant proportion of major ions [91, 92]. Our results also concur with previous work that showed marine bacteria have distinct regulatory networks for environmental sensing [11]. Collectively, these results identify environmental sensing capacity as a key genomic trait associated with bacterial lifestyles, ecological strategies, and environmental preferences.

### The sensor domain repertoire of bacterial metagenomes reveals consistent community responses to aridity

To evaluate whether environmental sensing capacity reflects environmental adaptation at the community level, we analyzed sensor domain composition across 255 soil metagenomes spanning a broad aridity gradient (Fig. 4A). Sensor domain composition was significantly associated with aridity (Mantel test r = 0.10, P = 0.008; Fig. S9), and unsupervised clustering based solely on sensor repertoires clearly separated metagenomes from arid and humid sites (Fig. S10). Likewise, db-RDA identified a distinct separation between arid and humid communities, with soil pH, conductivity, soil organic carbon, and net primary productivity explaining much of this variation (Fig. 4B; Table S3). As reported previously for deep subsurface microbial communities, variation in environmental conditions can be reflected in community-level sensor repertoires [70]. These findings align with previous evidence that aridity shapes soil microbial communities by selecting for taxa able to cope with the multiple environmental stressors (e.g. desiccation, UV radiation, osmotic stress) typical of arid soils [93, 94]. Beyond predicting distinct ecosystems [12], sensor domain repertoires can thus also be leveraged to distinguish bacterial communities with distinct environmental preferences within the same environment.

**Figure 4.**
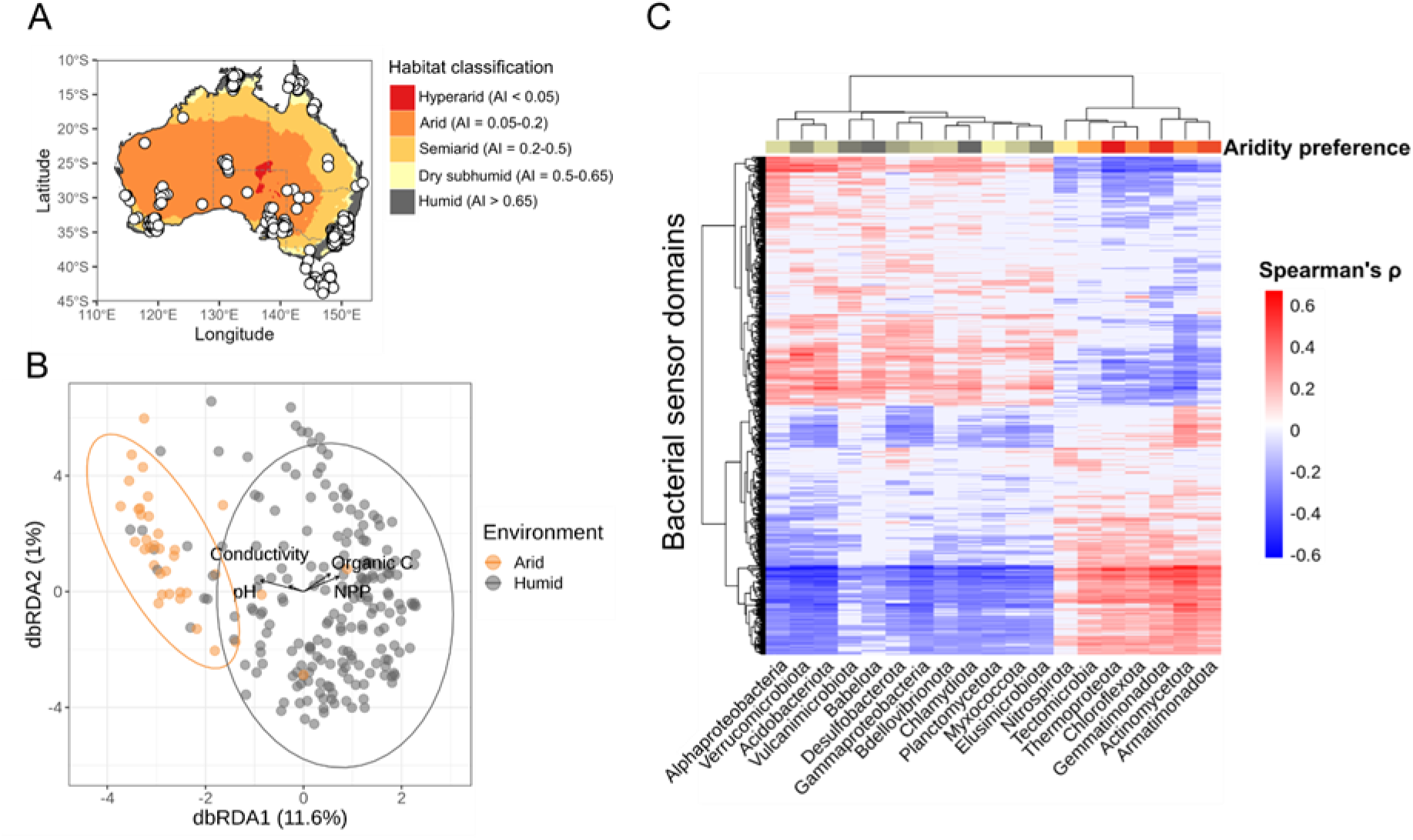
Associations between the composition of bacterial sensor domains and aridity across soil metagenomes. (A) Geographical distribution and aridity of sites where the metagenomic samples included in the study were collected. (B) Distance-based redundancy analysis (db-RDA) of the composition of sensor domains across metagenomes and the statistically significant associations with environmental factors. Differences in the sensor domain composition of metagenomes were based on Bray-Curtis distances (N = 255 metagenomes). (C) Heatmap showing significant Spearman correlations between the relative abundance of individual phyla and individual sensory domains. Rows and columns have been reordered following a complete hierarchical clustering of Euclidean distances. Correlations include FDR p-value correction. Column annotation shows the aridity preference index of the individual phyla which indicates the relative abundance change of phyla in samples from arid (AI < 0.65) over humid sites (AI ≥ 0.65).

We next examined whether these patterns were linked to taxonomic turnover along the gradient. Phyla enriched in arid soils included Actinomycetota, Chloroflexota and Gemmatimonadota, whereas Acidobacteriota, Pseudomonadota and Verrucomicrobiota were more abundant in humid sites (Fig. 4C). Correlations between phylum abundances and individual sensor domains revealed distinct clusters of domains associated with bacterial groups preferring arid or humid conditions. In total, 1,620 sensor domains were associated with aridity-associated phyla and 1,621 with phyla preferring humid conditions (Supplementary Dataset S3). Using a Support Vector Machine (SVM) classifier, we elucidated which sensor domains were most strongly associated with arid or humid conditions, identifying 100 sensor domains associated with arid conditions and 127 associated with humid conditions (Supplementary Dataset S3). Additionally, both groups of selected sensors showed a wide phylogenetic distribution, highlighting that despite the strong phylogenetic signal observed in the composition of sensor domains a large portion of these domains is not restricted to particular clades (Supplementary Fig. S12). Given the strong phylogenetic signal observed previously (Fig. 2A; Supplementary Fig. S1), the large number of sensors that occur across a broad range of phyla with shared environmental preference strongly suggests these likely underlie a common role in aridity adaptation.

### Identification of ligand binding motifs (LBMs) associated with bacterial communities from arid sites

We finally investigated associations between specific LBMs and aridity to determine whether environmental variation is reflected in sensory-domain function. We therefore analyzed the composition of sensor domains carrying experimentally characterized LBMs across soil metagenomes spanning an aridity gradient (Fig. 5A). Overall, the most widespread motifs were those binding divalent cations (96.1% of metagenomes) and 4-hydroxycinnamic acid (84.3%), whereas motifs binding carboxylates (18.8%) and b-type heme (16.9%) were the least prevalent (Fig. 5A). Similar to the analysis using full sensor domain sequences, LBM composition mostly separated metagenomes from arid sites from the rest (Fig. 5A), indicating that environmental preferences are associated not only with differences in sensory repertoires but also with variation in the environmental cues that bacterial communities can perceive.

**Figure 5.**
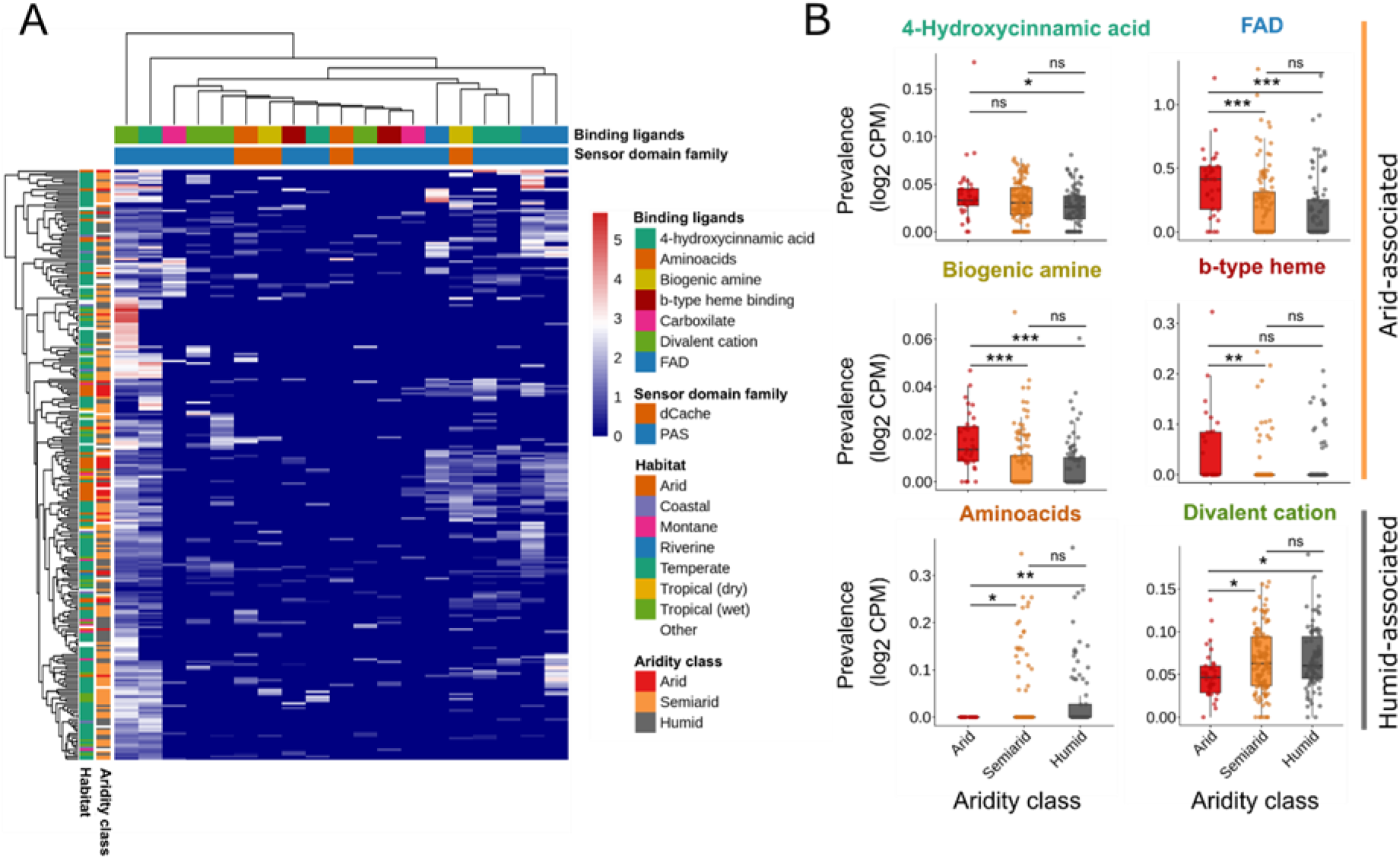
Associations between the prevalence of experimentally described ligand binding motifs and aridity in soil bacterial metagenomes. (A) Heatmap showing the prevalence of experimentally characterized ligand binding motifs across sensory domains present in the soil metagenomes across an aridity gradient. The prevalence of ligand binding motifs is expressed in counts per million (log_2_CPM). Metagenomes were classified based on their habitats of origin and the Aridity Index (AI) of their geographical location (row annotations). Column annotations indicate the protein family and characterized ligand of each sensor domain. (B) Prevalence in counts per million reads of experimentally characterized ligand binding motifs from sensor domains found in bacterial metagenomes from arid (AI ≤ 0.2), semiarid (AI = 0.2-0.65), and humid sites (AI ≥ 0.65). Statistical significance based on Mann Whitney U tests set at P < 0.05*, P < 0.01**, or P < 0.001***. N_Arid_ = 33; N_Semiarid_ = 117; N_Humid_ = 86.

Several LBMs were significantly enriched in metagenomes from arid soils, including motifs binding FAD (Kruskal-Wallis χ²(2) = 22.39, P < 0.001), biogenic amines (χ²(2) = 28.76, P < 0.001), b-type heme groups (χ²(2) = 10.19, P = 0.006), and 4-hydroxycinnamic acid (χ²(2) = 7.07, P = 0.029) (Fig. 5B). In contrast, motifs binding divalent cations (χ²(2) = 8.06, P = 0.018) and amino acids (χ²(2) = 10.12, P = 0.006) were more prevalent in humid soils (Fig. 5B). No significant differences were observed for carboxylate- or c-type-heme-binding motifs (χ²(2) = 4.64, P = 0.09; Supplementary Fig. S13). These results indicate that adaptation to arid environments likely involves not only increased tolerance to environmental stress but also distinct capacities to perceive environmentally relevant signals.

The enrichment of FAD- and b-type-heme-binding domains in arid soils is consistent with a greater importance of redox and oxygen sensing under water limitation, where respiration, oxidative stress, and energy metabolism can fluctuate substantially [95–98]. Likewise, the higher prevalence of biogenic-amine-binding motifs is consistent with the role of compounds such as choline and glycine betaine as osmoprotectants under drought stress [97, 99]. In contrast, humid soils were enriched in amino-acid- and divalent-cation-binding motifs. Amino acids function both as nutrients and environmental cues involved in habitat selection, colonization, and resource acquisition [23], and their enrichment may reflect the greater substrate availability and microbial activity characteristic of wetter soils. Similarly, the higher prevalence of divalent-cation-binding domains may be linked to enhanced mineral weathering and ion mobilization in humid environments [100]. Although mechanistic knowledge of bacterial sensory functions remains limited, these results demonstrate that currently characterized LBMs can generate testable hypotheses linking environmental sensing to ecological adaptation. The sensor domain sequences and LBMs identified here could help track microbial responses to aridification and drought [101], inform the design of climate-resilient microbiomes through sensory traits associated with environmental adaptation [102], and support the development of biosensors based on naturally occurring microbial sensory modules [103–105].

## Conclusions

Microbial traits are commonly organized along major ecological axes associated with resource acquisition and environmental response strategies [106–108]. We showed that environmental sensing capacity is associated with core evolutionary processes, microbial lifestyles, ecological strategies, and environmental preferences, so environmental sensing capacity is a trait closely linked to these ecological axes [109]. For example, nutrient-rich and heterogeneous habitats that favor copiotrophic lifestyles also select for more diverse sensing repertoires [110], while stress-tolerant and generalist organisms benefit from enhanced capacities to perceive and respond to environmental change [111]. Environmental sensing capacity is currently distributed across KEGG Environmental Information Processing pathways [112], COG category T (signal transduction mechanisms, [113], and multiple SEED regulation and signaling subsystems [114], but is not represented as a unified functional category in any of the major annotation frameworks. This fragmentation may have contributed to environmental sensing being overlooked as a coherent ecological trait. We propose that environmental sensing capacity constitutes an overlooked axis of microbial trait variation and should be incorporated into trait-based frameworks aimed at understanding microbial life histories and responses to environmental change. To more comprehensively describe environmental sensing, similar studies with other receptor families, such as chemoreceptors and receptors that control second messenger levels are needed [115]. These receptors share fundamentally the same set of sensor domains and there is currently a major research need to establish to which degree the different sensory families are interconnected [20]. Description of the signal transduction pathways connecting sensors with response genes would establish the missing mechanistic link between perception and adaptive response in bacteria.

## Acknowledgements

J.R. acknowledges funding from the Spanish Ministry for Science, Innovation and Universities/Agencia Estatal de Investigación (MicroDroughtPredict PID2023-151209NA-I00 and Ramón y Cajal Fellowship RYC2024-049086-I), and CSIC (Proyectos Intramurales Especiales, PREDEX 20263AT004) awarded to J.R. A.P.-G. acknowledges funding from a Ramón y Cajal Fellowship (RyC2021-032424-I) from the Spanish Ministry for Science, Innovation and Universities/Agencia Estatal de Investigación and PRTR (EU) awarded to A.P.-G. J.R. and A.P.-G. acknowledge support from the Scientific Network BCB funded by CSIC (Spain). T.K. acknowledges was funded by the Spanish Ministry for Science, Innovation and Universities/Agencia Estatal de Investigación 10.13039/501100011033 grant PID2023-146216NB-I00 awarded to T.K. R.S.A acknowledges funding from the JAE Intro programme 2025 from CSIC.

## Author contributions

J.R. conceptualized, designed, and coordinated the study. R.S.A. performed the analyses with assistance from A.C.-P. and A.F.-V. T.K. and A.P.-G. provided critical intellectual input and made substantial edits to the final text. R.S.A., J.R. and G.S. wrote the manuscript with input from all co-authors.

## References

1. Alvarez AF, Georgellis D. Environmental adaptation and diversification of bacterial two-component systems. Curr Opin Microbiol 2023;76:102399.

2. Galperin MY. What bacteria want. Environ Microbiol 2018;20:4221–4229.

3. Ulrich LE, Koonin E V, Zhulin IB. One-component systems dominate signal transduction in prokaryotes. Trends Microbiol 2005;13:52–56.

4. Parkinson JS. Signal transduction schemes of bacteria. Cell 1993;73:857–871.

5. Zhulin IB, Nikolskaya AN, Galperin MY. Common extracellular sensory domains in transmembrane receptors for diverse signal transduction pathways in bacteria and archaea. J Bacteriol 2003;185:285–294.

6. Lacal J et al. Sensing of environmental signals: classification of chemoreceptors according to the size of their ligand binding regions. Environ Microbiol 2010;12:2873–2884.

7. Galperin MY. A census of membrane-bound and intracellular signal transduction proteins in bacteria: bacterial IQ, extroverts and introverts. BMC Microbiol 2005;5:35.

8. Alm E, Huang K, Arkin A. The evolution of two-component systems in bacteria reveals different strategies for niche adaptation. PLoS Comput Biol 2006;2:e143.

9. Bhar S, Bose T, Mande SS. Can pathogenic and nonpathogenic bacteria be distinguished by sensory protein abundance? Appl Environ Microbiol 2020;86:e00478–20.

10. Rodrigue A et al. Cell signalling by oligosaccharides. Two-component systems in Pseudomonas aeruginosa: why so many? Trends Microbiol 2000;8:498–504.

11. Held NA et al. Unique patterns and biogeochemical relevance of two-component sensing in marine bacteria. MSystems 2019;4:10–1128.

12. Park H et al. A bacterial sensor taxonomy across earth ecosystems for machine learning applications. Msystems 2024;9:e00026-23.

13. Guan N et al. Microbial response to environmental stresses: from fundamental mechanisms to practical applications. Appl Microbiol Biotechnol 2017;101:3991–4008.

14. Krell T et al. Bacterial sensor kinases: diversity in the recognition of environmental signals. Annu Rev Microbiol 2010;64:539–559.

15. Hoch JA. Two-component and phosphorelay signal transduction. Curr Opin Microbiol 2000;3:165–170.

16. Mascher T, Helmann JD, Unden G. Stimulus perception in bacterial signal-transducing histidine kinases. Microbiol Mol Biol Rev 2006;70:910–938.

17. Galperin MY. Diversity of structure and function of response regulator output domains. Curr Opin Microbiol 2010;13:150–159.

18. Cerna-Vargas JP et al. Amine-recognizing domain in diverse receptors from bacteria and archaea evolved from the universal amino acid sensor. Proc Natl Acad Sci 2023;120:e2305837120.

19. Fernández M et al. Identification of a chemoreceptor that specifically mediates chemotaxis toward metabolizable purine derivatives. Mol Microbiol 2016;99:34–42.

20. Sanchis-López C et al. Prevalence and specificity of chemoreceptor profiles in plant-associated bacteria. Msystems 2021;6:10–1128.

21. Matilla MA et al. Structural and functional diversity of sensor domains in bacterial transmembrane receptors. Trends Microbiol 2025.

22. Gavira JA et al. How bacterial chemoreceptors evolve novel ligand specificities. Mbio 2020;11:10–1128.

23. Matilla MA, Krell T. Bacterial amino acid chemotaxis: a widespread strategy with multiple physiological and ecological roles. J Bacteriol 2024;206:e00300–24.

24. Henry JT, Crosson S. Ligand-binding PAS domains in a genomic, cellular, and structural context. Annu Rev Microbiol 2011;65:261–286.

25. Taylor BL, Rebbapragada A, Johnson MS. The FAD-PAS domain as a sensor for behavioral responses in Escherichia coli. Antioxid Redox Signal 2001;3:867–879.

26. Gumerov VM et al. Amino acid sensor conserved from bacteria to humans. Proc Natl Acad Sci 2022;119:e2110415119.

27. Upadhyay AA et al. Cache domains that are homologous to, but different from PAS domains comprise the largest superfamily of extracellular sensors in prokaryotes. PLoS Comput Biol 2016;12:e1004862.

28. Day CJ et al. A direct-sensing galactose chemoreceptor recently evolved in invasive strains of Campylobacter jejuni. Nat Commun 2016;7:13206.

29. Boyeldieu A et al. Combining two optimized and affordable methods to assign chemoreceptors to a specific signal. Anal Biochem 2021;620:114139.

30. Wu R et al. Insight into the sporulation phosphorelay: crystal structure of the sensor domain of Bacillus subtilis histidine kinase, KinD. Protein Sci 2013;22:564–576.

31. Zhang L et al. Sensing of autoinducer-2 by functionally distinct receptors in prokaryotes. Nat Commun 2020;11:5371.

32. Monteagudo-Cascales E et al. Ubiquitous purine sensor modulates diverse signal transduction pathways in bacteria. Nat Commun 2024;15:5867.

33. Webb BA et al. Sinorhizobium meliloti chemotaxis to quaternary ammonium compounds is mediated by the chemoreceptor McpX. Mol Microbiol 2017;103:333–346.

34. Corral-Lugo A et al. High-affinity chemotaxis to histamine mediated by the TlpQ chemoreceptor of the human pathogen Pseudomonas aeruginosa. MBio 2018;9:10–1128.

35. Sleator RD, Hill C. Bacterial osmoadaptation: the role of osmolytes in bacterial stress and virulence. FEMS Microbiol Rev 2002;26:49–71.

36. Zaprasis A et al. Uptake of amino acids and their metabolic conversion into the compatible solute proline confers osmoprotection to Bacillus subtilis. Appl Environ Microbiol 2015;81:250–259.

37. Gilabert MJ et al. Acetylcholine signaling regulates osmotic stress adaptation in the phytopathogen Dickeya solani. Microbiol Res 2026;128646.

38. Shen J et al. Spatial Variability of Microbial Communities and Salt Distributions Across a Latitudinal Aridity Gradient in the Atacama Desert: Shen J. et al. Microb Ecol 2021;82:442–458.

39. Coleine C et al. Dryland microbiomes reveal community adaptations to desertification and climate change. ISME J 2024;18:wrae056.

40. Krogh A et al. Predicting transmembrane protein topology with a hidden Markov model: application to complete genomes. J Mol Biol 2001;305:567–580.

41. Xing J, Gumerov VM, Zhulin IB. Origin and functional diversification of PAS domain, a ubiquitous intracellular sensor. Sci Adv 2023;9:eadi4517.

42. Potter SC, et al. HMMER web server: 2018 update. Nucleic Acids Res 2018;46:W200–W204.

43. Mistry J et al. Pfam: The protein families database in 2021. Nucleic Acids Res 2021;49:D412–D419.

44. Parks DH et al. GTDB: an ongoing census of bacterial and archaeal diversity through a phylogenetically consistent, rank normalized and complete genome-based taxonomy. Nucleic Acids Res 2022;50:D785–D794.

45. Parks DH et al. CheckM: assessing the quality of microbial genomes recovered from isolates, single cells, and metagenomes. Genome Res 2015;25:1043–1055.

46. Bissett A et al. Introducing BASE: the Biomes of Australian Soil Environments soil microbial diversity database. Gigascience 2016;5:s13742–016.

47. Lebre PH, De Maayer P, Cowan DA. Xerotolerant bacteria: surviving through a dry spell. Nat Rev Microbiol 2017;15:285–296.

48. Zomer RJ, Xu J, Trabucco A. Version 3 of the global aridity index and potential evapotranspiration database. Sci Data 2022;9:409.

49. Andrews S. FastQC: a quality control tool for high throughput sequence data. 2010. 2017.

50. Li D et al. MEGAHIT: an ultra-fast single-node solution for large and complex metagenomics assembly via succinct de Bruijn graph. Bioinformatics 2015;31:1674–1676.

51. Gurevich A et al. QUAST: quality assessment tool for genome assemblies. Bioinformatics 2013;29:1072–1075.

52. Woodcroft BJ et al. Comprehensive taxonomic identification of microbial species in metagenomic data using SingleM and Sandpiper. Nat Biotechnol 2026;44:948–953.

53. Pardoux R, Dolla A, Aubert C. Metal-containing PAS/GAF domains in bacterial sensors. Coord Chem Rev 2021;442:214000.

54. Podlesny D et al. metaTraits: a large-scale integration of microbial phenotypic trait information. Nucleic Acids Res 2026;54:D835–D841.

55. Kim CY et al. Planetary microbiome structure and generalist-driven gene flow across disparate habitats. Cell 2026;189:2073–2091.

56. Dixon P. VEGAN, a package of R functions for community ecology. J Veg Sci 2003;14:927–930.

57. Fritz SA, Purvis A. Phylogenetic diversity does not capture body size variation at risk in the world’s mammals. Proc R Soc B Biol Sci 2010;277:2435.

58. Paradis E, Schliep K. ape 5.0: an environment for modern phylogenetics and evolutionary analyses in R. Bioinformatics 2019;35:526–528.

59. Letunic I, Bork P. Interactive Tree Of Life (iTOL) v5: an online tool for phylogenetic tree display and annotation. Nucleic Acids Res 2021;49:W293–W296.

60. Lin Z et al. Evolutionary-scale prediction of atomic-level protein structure with a language model. Science (80-) 2023;379:1123–1130.

61. Ishii E, Eguchi Y. Diversity in sensing and signaling of bacterial sensor histidine kinases. Biomolecules 2021;11:1524.

62. Kim D, Forst S. Genomic analysis of the histidine kinase family in bacteria and archaea. Microbiology 2001;147:1197–1212.

63. Galperin MY. Structural classification of bacterial response regulators: diversity of output domains and domain combinations. J Bacteriol 2006;188:4169–4182.

64. Matilla MA et al. A catalogue of signal molecules that interact with sensor kinases, chemoreceptors and transcriptional regulators. FEMS Microbiol Rev 2022;46:fuab043.

65. Koretke KK et al. Evolution of two-component signal transduction. Mol Biol Evol 2000;17:1956–1970.

66. Murphy CL et al. Genomic characterization of three novel Desulfobacterota classes expand the metabolic and phylogenetic diversity of the phylum. Environ Microbiol 2021;23:4326–4343.

67. Beck C et al. The diversity of cyanobacterial metabolism: genome analysis of multiple phototrophic microorganisms. BMC Genomics 2012;13:56.

68. Li S-H, Kang I, Cho J-C. Metabolic versatility of the family Halieaceae revealed by the genomics of novel cultured isolates. Microbiol Spectr 2023;11:e03879–22.

69. Li L et al. Globally distributed Myxococcota with photosynthesis gene clusters illuminate the origin and evolution of a potentially chimeric lifestyle. Nat Commun 2023;14:6450.

70. Goldman AL et al. Microbial sensor variation across biogeochemical conditions in the terrestrial deep subsurface. Msystems 2024;9:e00966–23.

71. Chen M-Y et al. Comparative genomics reveals insights into cyanobacterial evolution and habitat adaptation. ISME J 2021;15:211–227.

72. Sorensen JW et al. Ecological selection for small microbial genomes along a temperate-to-thermal soil gradient. Nat Microbiol 2019;4:55–61.

73. Giovannoni SJ et al. Genome streamlining in a cosmopolitan oceanic bacterium. Science (80-) 2005;309:1242–1245.

74. Yus E et al. Impact of genome reduction on bacterial metabolism and its regulation. Science (80-) 2009;326:1263–1268.

75. Genova R et al. Chemotaxis to plant defense compounds in phytopathogens. Plos Pathog 2026;22:e1014240.

76. Capra EJ, Laub MT. Evolution of two-component signal transduction systems. Annu Rev Microbiol 2012;66:325–347.

77. D’Souza G et al. Less is more: selective advantages can explain the prevalent loss of biosynthetic genes in bacteria. Evolution (N Y) 2014;68:2559–2570.

78. Puente-Sánchez F et al. Cross-biome microbial networks reveal functional redundancy and suggest genome reduction through functional complementarity. Commun Biol 2024;7:1046.

79. Ramoneda J et al. Taxonomic and environmental distribution of bacterial amino acid auxotrophies. Nat Commun 2023;14:7608.

80. Morris JJ, Lenski RE, Zinser ER. The Black Queen Hypothesis: evolution of dependencies through adaptive gene loss. MBio 2012;3:10–1128.

81. Nielsen DA et al. Aerobic bacteria and archaea tend to have larger and more versatile genomes. Oikos 2021;130:501–511.

82. Barberán A et al. Why are some microbes more ubiquitous than others? Predicting the habitat breadth of soil bacteria. Ecol Lett 2014;17:794–802.

83. Maschmann ZA et al. Redox properties and PAS domain structure of the Escherichia coli energy sensor Aer indicate a multistate sensing mechanism. J Biol Chem 2022;298.

84. Flamholz AI et al. Annotation-free prediction of microbial dioxygen utilization. Msystems 2024;9:e00763–24.

85. Jabłońska J, Tawfik DS. The number and type of oxygen-utilizing enzymes indicates aerobic vs. anaerobic phenotype. Free Radic Biol Med 2019;140:84–92.

86. Bueno de Mesquita CP, Stallard-Olivera E, Fierer N. Predicting oxygen levels in microbial habitats using a metagenome-based approach. Msystems 2026;e00545–26.

87. Mitchell JG, Okubo A, Fuhrman JA. Microzones surrounding phytoplankton form the basis for a stratified marine microbial ecosystem. Nature 1985;316:58–59.

88. Stocker R et al. Rapid chemotactic response enables marine bacteria to exploit ephemeral microscale nutrient patches. Proc Natl Acad Sci 2008;105:4209–4214.

89. Smriga S et al. Chemotaxis toward phytoplankton drives organic matter partitioning among marine bacteria. Proc Natl Acad Sci 2016;113:1576–1581.

90. Likens GE. Biogeochemistry of inland waters. Academic press, 2010.

91. Pilson MEQ. An Introduction to the Chemistry of the Sea. 1998.

92. Lebrato M et al. Global variability in seawater Mg: Ca and Sr: Ca ratios in the modern ocean. Proc Natl Acad Sci 2020;117:22281–22292.

93. Neilson JW et al. Significant impacts of increasing aridity on the arid soil microbiome. MSystems 2017;2:10–1128.

94. Li C et al. The adjustment of life history strategies drives the ecological adaptations of soil microbiota to aridity. Mol Ecol 2022;31:2920–2934.

95. Ramírez-Flandes S, González B, Ulloa O. Redox traits characterize the organization of global microbial communities. Proc Natl Acad Sci 2019;116:3630–3635.

96. Farhana A et al. Environmental heme-based sensor proteins: implications for understanding bacterial pathogenesis. Antioxid Redox Signal 2012;17:1232–1245.

97. Schimel JP. Life in dry soils: effects of drought on soil microbial communities and processes. Annu Rev Ecol Evol Syst 2018;49:409–432.

98. Bouskill NJ et al. Climate history modulates stress responses of common soil bacteria under experimental drought. ISME J 2025;19:wraf075.

99. Wood JM. Bacterial osmoregulation: a paradigm for the study of cellular homeostasis. Annu Rev Microbiol 2011;65:215–238.

100. Fang Q et al. Mineral weathering is linked to microbial priming in the critical zone. Nat Commun 2023;14:345.

101. Knight CG et al. Soil microbiomes show consistent and predictable responses to extreme events. Nature 2024;636:690–696.

102. Silverstein MR, Segrè D, Bhatnagar JM. Environmental microbiome engineering for the mitigation of climate change. Glob Chang Biol 2023;29:2050–2066.

103. Gui Q et al. The application of whole cell-based biosensors for use in environmental analysis and in medical diagnostics. Sensors 2017;17:1623.

104. Moraskie M et al. Microbial whole-cell biosensors: Current applications, challenges, and future perspectives. Biosens Bioelectron 2021;191:113359.

105. Fu Y et al. Development of a two component system based biosensor with high sensitivity for the detection of copper ions. Commun Biol 2024;7:1407.

106. Ho A, Di Lonardo DP, Bodelier PLE. Revisiting life strategy concepts in environmental microbial ecology. FEMS Microbiol Ecol 2017;93:fix006.

107. Malik AA et al. Defining trait-based microbial strategies with consequences for soil carbon cycling under climate change. ISME J 2020;14:1–9.

108. Stone BWG et al. Life history strategies among soil bacteria—dichotomy for few, continuum for many. ISME J 2023;17:611–619.

109. Westoby M et al. Trait dimensions in bacteria and archaea compared to vascular plants. Ecol Lett 2021;24:1487–1504.

110. Lauro FM et al. The genomic basis of trophic strategy in marine bacteria. Proc Natl Acad Sci 2009;106:15527–15533.

111. López-Maury L, Marguerat S, Bähler J. Tuning gene expression to changing environments: from rapid responses to evolutionary adaptation. Nat Rev Genet 2008;9:583–593.

112. Kanehisa M, Goto S. KEGG: kyoto encyclopedia of genes and genomes. Nucleic Acids Res 2000;28:27–30.

113. Galperin MY, et al. COG database update 2024. Nucleic Acids Res 2025;53:D356–D363.

114. Overbeek R et al. The subsystems approach to genome annotation and its use in the project to annotate 1000 genomes. Nucleic Acids Res 2005;33:5691–5702.

115. Velando F et al. Chemoreceptor family in plant-associated bacteria responds preferentially to the plant signal molecule glycerol 3-phosphate. Genome Biol 2025;26:260.

