## Supplementary for "Environmental sensing capacity predicts bacterial ecological strategies and environmental preferences"

Roman Sanz-Altxu et al.

This file includes:

Figs. S1 to S13

Tables S1 and S4

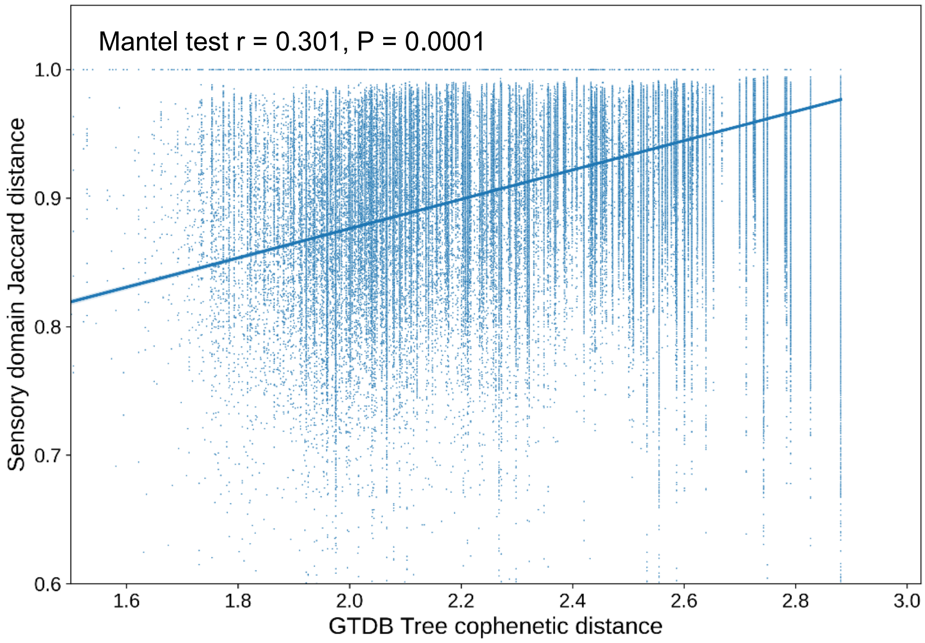


**Supplementary Figure S1.** Association between differences in the sensor domain repertoires of bacteria and their phylogenetic distances. Cophenetic distances were obtained from the Genome Taxonomy Database (GTDB) phylogeny.

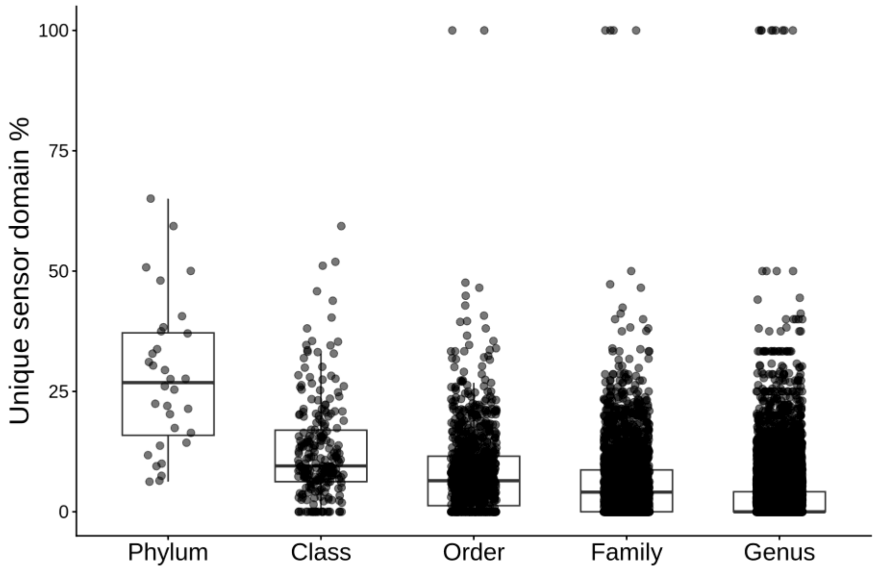


**Supplementary Figure S2.** Distribution of unique bacterial sensor domains per taxonomic level.

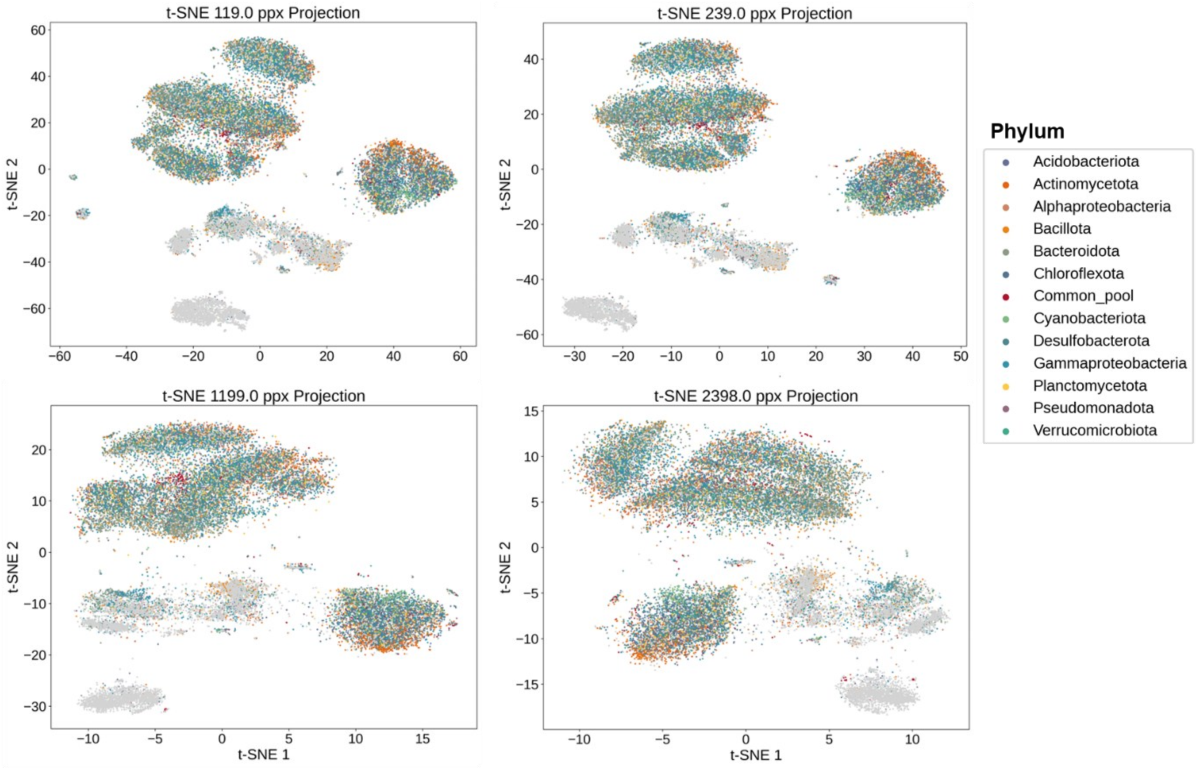


**Supplementary Figure S3. Visualization of extracytosolic sensor domain sequence diversity across bacterial phyla.** Each point represents an individual sensor domain sequence, positioned by t-distributed stochastic neighbor embedding (t-SNE) of pairwise sequence identity. Points are colored according to whether the corresponding sequence is restricted to a single phylum (phylum-specific) or shared across multiple phyla (part of a common sequence pool), as indicated in the legend. Panels show the same embedding computed at different perplexity values (0.5%, 1%, 5% and 10% of our dataset size), a t-SNE parameter that approximates the effective number of nearest neighbors considered when preserving local structure; lower perplexity emphasizes fine-scale, local clustering, while higher perplexity captures broader, more global relationships between sequences. Clustering patterns reflect underlying sequence similarity, with tighter groupings indicating higher local sequence conservation.

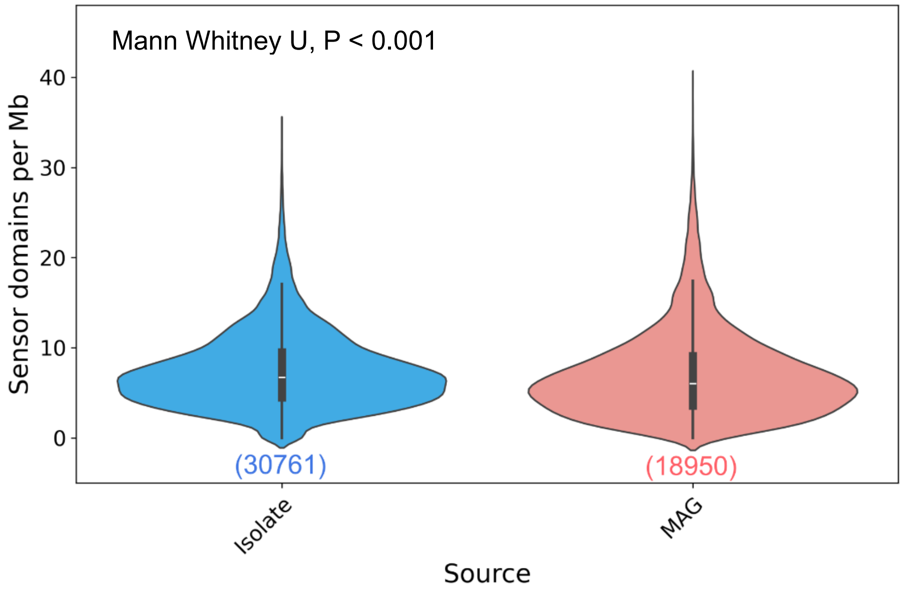


**Supplementary Figure S4.** Prevalence of extracytosolic and PAS sensor domains in genomes coming from bacterial isolates and metagenome-assembled genomes (MAGs).

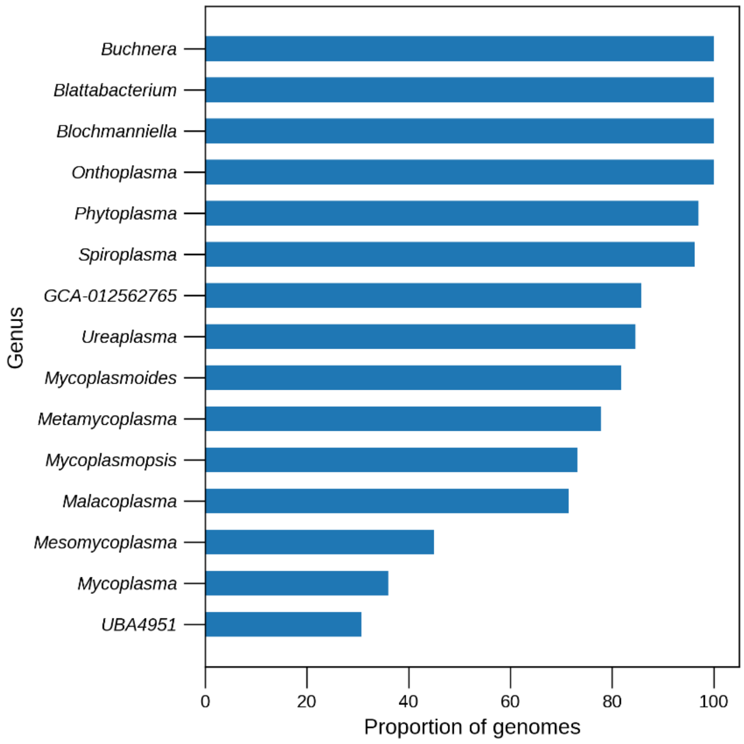


**Supplementary Figure S5.** Proportion of genomes that did not contain any sensor domain sequences, grouped (when known) by their genus affiliation. These genera belong to either endosymbionts (*Buchnera, Blattabacteria,* and *Blochmanniella*), plant and insect parasites (*Onthoplasma, Phytoplasma,* and *Spiroplasma*), or animal-associated parasites and commensals (*Ureaplasma, Mycoplasmoides, Metamycoplasma, Mycoplasmopsis, Malacoplasma., Mesomycoplasma*, and *Mycoplasma*).

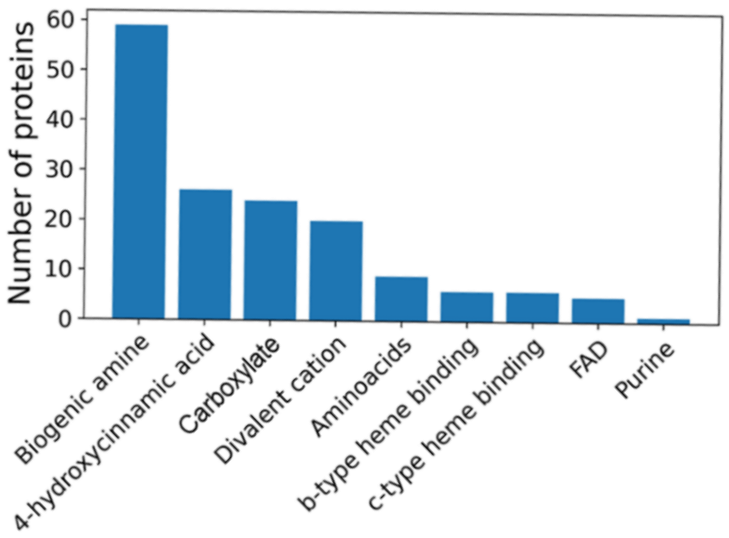


**Supplementary Figure S6.** Number of unique sensor domains containing ligand binding motifs (LBMs) for different ligands within our extracytosolic and PAS sensor domain database.

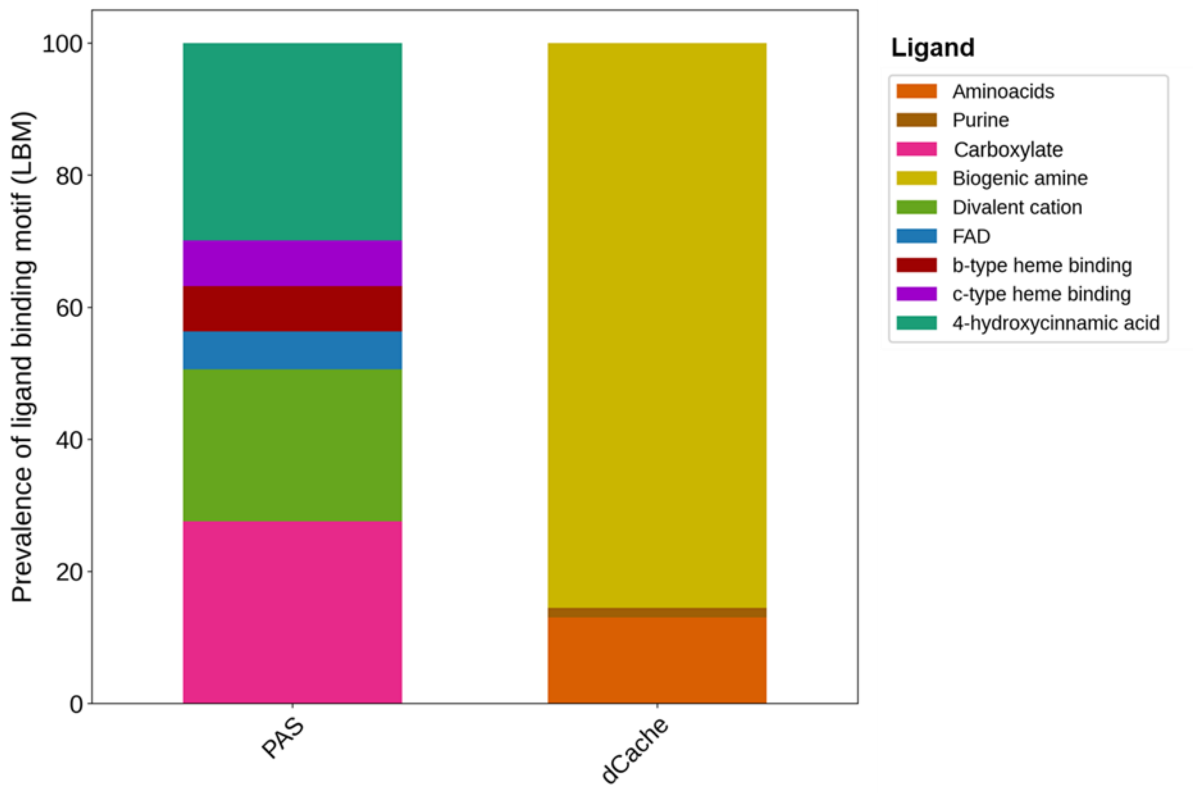


**Supplementary Figure S7.** Proportions of ligand binding motifs (LBMs) identified across the two main sensor domain families investigated here.

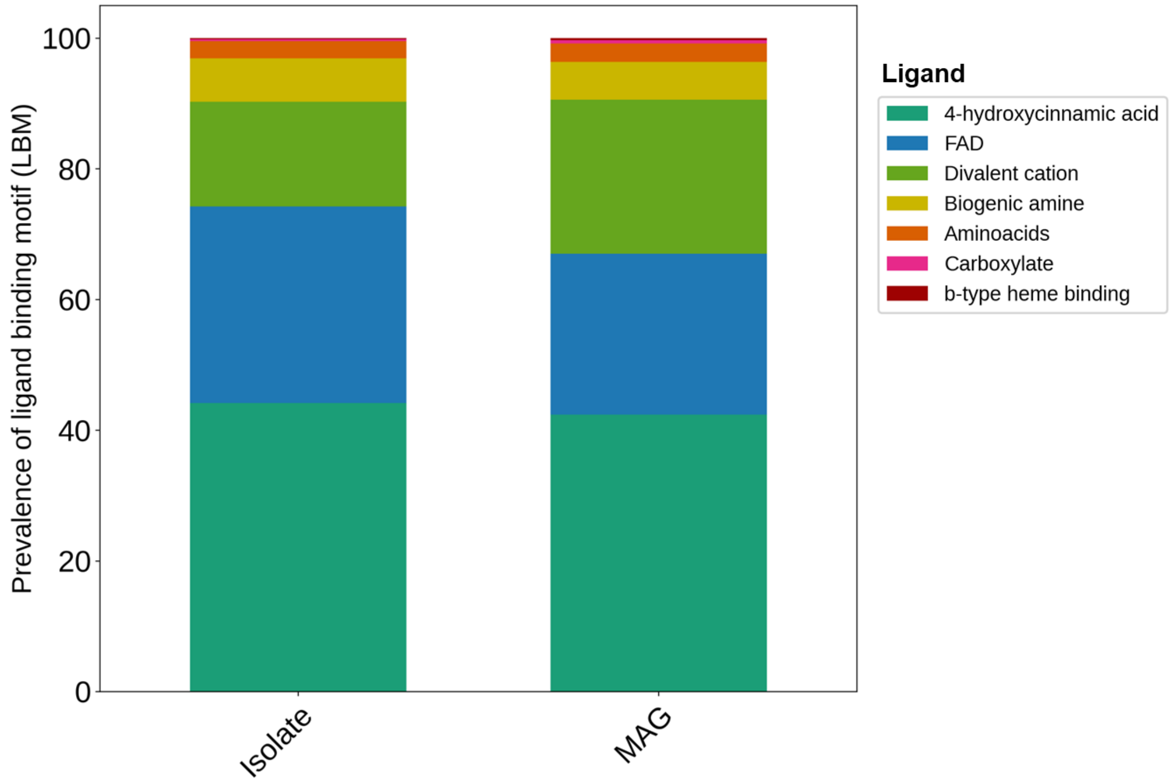


**Supplementary Figure S8.** Proportions of ligand binding motifs (LBMs) identified in our sensor domain database in genomes coming from bacterial isolates and metagenome-assembled genomes (MAGs). LBMs for carboxylate and c-type heme binding were excluded for being extremely underrepresented in genomes (<0.5% occurrence).

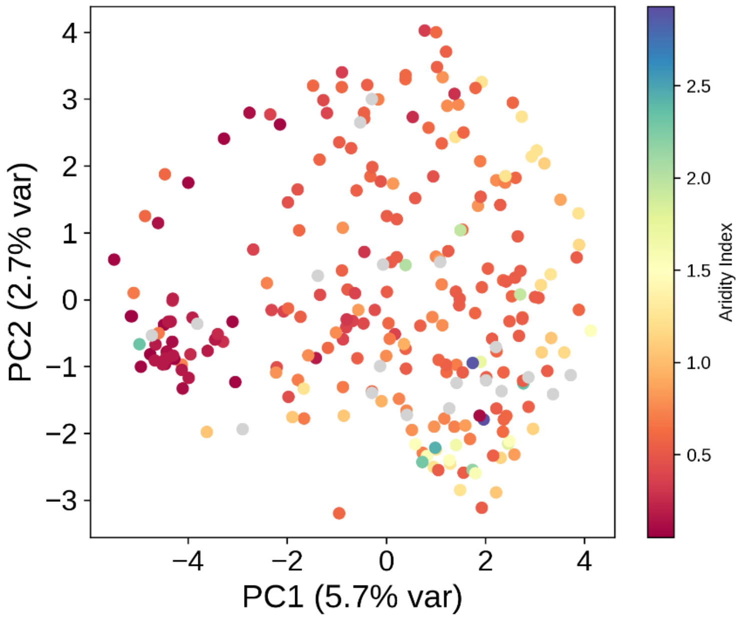


**Supplementary Figure S9.** Principal Component Analysis (PCA) of the sensory domain composition of soil bacterial metagenomes from an aridity gradient across the Australian continent (N = 255). Metagenomic data was obtained from Australian Microbiome Initiative Biomes of Australian Soil Environments (BASE) project [1].

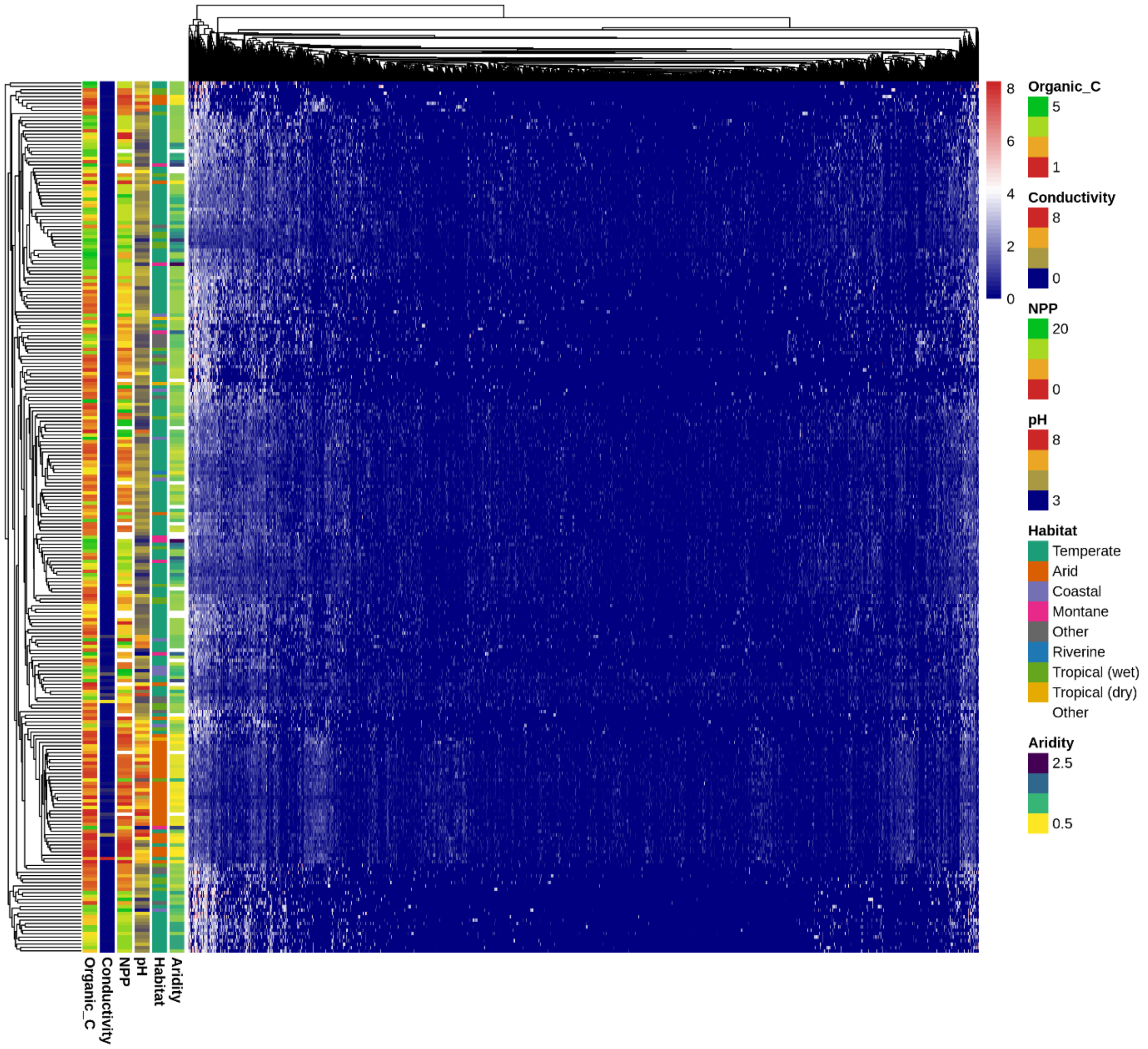


**Supplementary Figure S10.** Heatmap showing the sensory domain composition (columns) of soil bacterial metagenomes (rows) from an aridity gradient across Australia (N = 255). Rows and columns have been clustered following hierarchical clustering of their Euclidean distances.

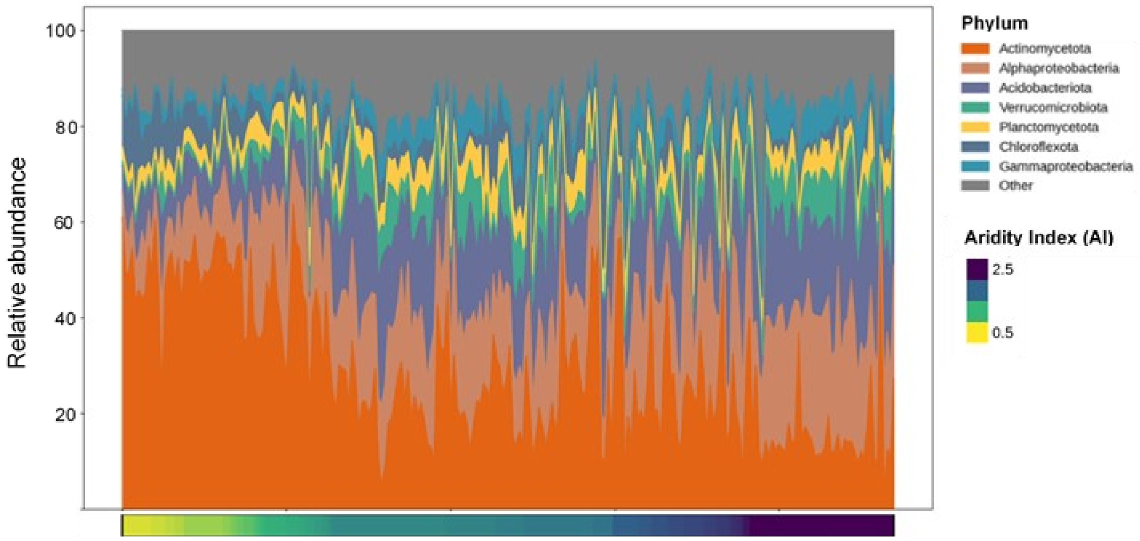


**Supplementary Figure S11.** Taxonomic profile of soil bacterial metagenomic samples from across Australia ordered by aridity (N = 255). Cumulative area values indicate relative abundance of represented phyla.

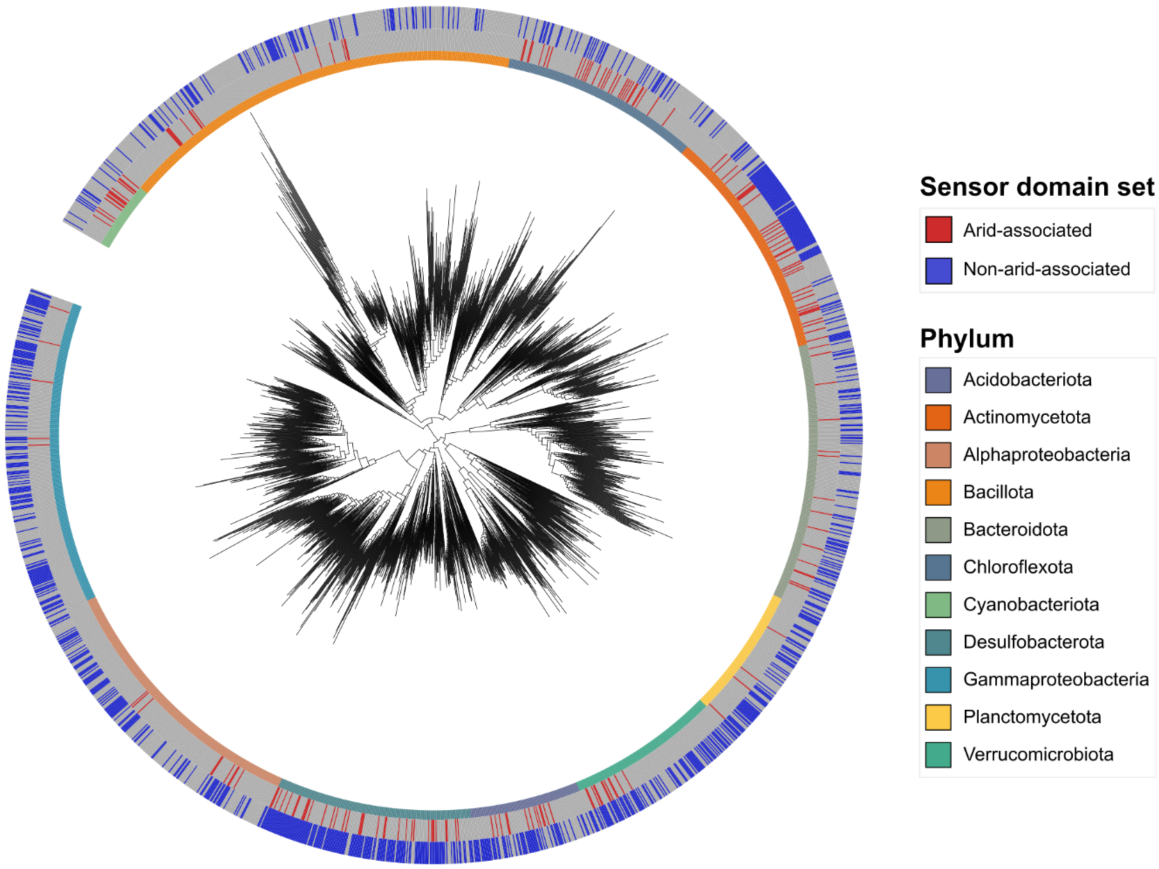


**Supplementary Figure S12.** Phylogenetic distribution of bacterial extracytosolic and PAS sensor domains identified as being indicative of soil communities from arid (N = 100) and humid sites (N = 127). The represented sensor domains had the highest discriminatory power in a Support Vector Machine (SVM) model describing differences in the sensor domain repertoires of bacterial communities from arid versus humid sites.

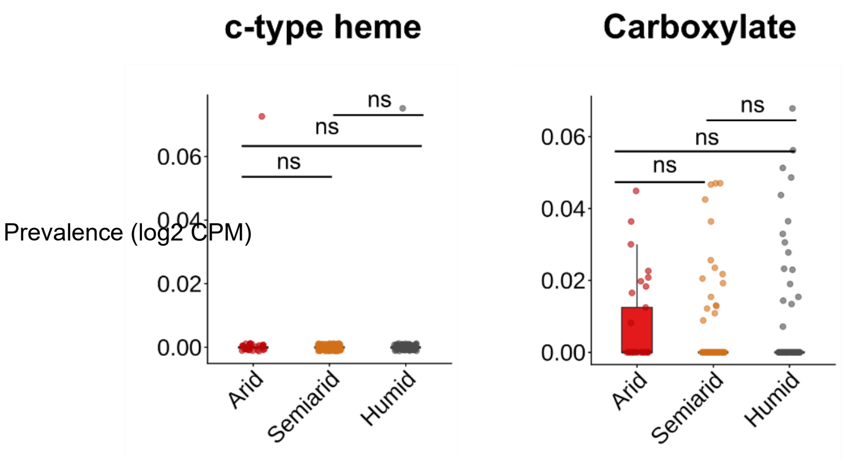


**Supplementary Figure S13.** Prevalence in counts per million reads of ligand binding motifs (LBMs) for binding c-type heme groups and carboxylate from sensor domains found in bacterial metagenomes from arid (aridity index AI ≤ 0.2), semiarid (AI = 0.2-0.65), and humid sites (AI ≥ 0.65). Statistical significance based on Mann Whitney U tests with Benjamini-Hochberg correction set at $P < 0.05$*, $P < 0.01$**, or $P < 0.001$***. $N_{Arid} = 33$; $N_{Semiarid} = 117$; $N_{Humid} = 86$.

**Supplementary Table S1.** Details of the sequences, ligands, and functions of ligand binding motifs (LBMs) that have been experimentally described in previous studies which we used here to determine associations between ligand-binding functions and bacterial taxa, ecological strategies, and environmental preferences.

| Motif | Family | Ligand | Function | Reference |
|---|---|---|---|---|
| [Y](4)[R](1)[W](13-17)[Y](27-34)[D] | dCache | Aminoacids | Chemotaxis for amino acids, some are involved in osmoregulation | [2] |
| [YWF](20-34)[WYF][WYF](20-25)[MLIVAC](1)[ST](17-21)[D] | dCache | Biogenic amine | Chemotaxis for compatible solutes, osmoregulation | [3] |
| [YF](7-8)[RKH](6-10)[WF][YF](21-28)[YF](1)[DN] | dCache | Purine | Chemotaxis, nutrient scavenging | [4] |
| C(7)C(1)C(17)C | GAF | Heme Group | Oxygen and redox sensing | [5] |
| N(1)R(2)Q(12)[R/K](13)[N](9)[N](18-20)[Q] | PAS | Flavin | Photosensing | [6] |
| [RSN](2)[NRS](5)[RK](6)[W](14)[N] | PAS | FAD | Oxygen and redox sensing | [6] |
| R(2)H(31-32)L(4)R | PAS | Carboxylate | Transport/Metabolism | [6] |
| R(1)Y(16)T(1)S(5)Y(17)K | PAS | Dicarboxylate | Transport/Metabolism | [6] |
| [ED](30)E(1)T(27)E | PAS | Divalent cation | Ion homeostasis | [6] |
| Y(3)E(22)C | PAS dCache | 4-hydroxycinnamic acid | Photosensing (PAS) and plant chemotaxis (dCache) | [6] |
| H(5)R(7)HI | PAS | b-type heme binding | Oxygen sensing | [6] |
| H(6)N(9)GM | PAS | b-type heme binding | Oxygen sensing | [6] |
| C(2)CH | PAS | c-type heme binding | Gas sensing | [6] |

**Supplementary Table S2.** Statistical summary of the tests to assess phylogenetic signal of ligand binding motif (LBM) composition in bacteria (N = 51343 genomes). The D phylogenetic index was obtained by treating the presence of a given LBM in genomes as a binary trait. Mantel test correlations were based on correlations between Jaccard distances of the LBM repertoires of bacterial genomes and cophenetic distances between those genomes within the Genome Taxonomy Database (GTDB) phylogeny.

| Ligand | Fritz and Purvis D | P-value | Mantel's r | P-value |
|---|---|---|---|---|
| Aminoacids | 0.670 | 0.014 | 0.005 | < 0.001 |
| Biogenic amine | 0.679 | < 0.001 | 0.015 | < 0.001 |
| b-type heme | 0.172 | 0.364 | -0.0001 | 0.526 |
| Divalent cation | 0.669 | < 0.001 | 0.0007 | 0.318 |
| FAD | 0.324 | < 0.001 | 0.019 | < 0.001 |
| 4-Hydroxycinnamic acid | 0.576 | < 0.001 | 0.017 | < 0.001 |

**Supplementary Table S3.** Statistical summary of co-occurrences of ligand binding motifs (LBMs) across bacterial genomes from the tree of life (N = 51343). Numbers indicate the FDR-corrected P-values obtained from pairwise Fisher's exact tests among ligands.

| Ligand | Aminoacids | Biogenic amine | b-type heme | Divalent cation | FAD | 4-Hydroxycinnamic acid |
|---|---|---|---|---|---|---|
| Aminoacids | | 0.085 | 0.86 | 0.214 | < 0.001 | < 0.001 |
| Biogenic amine | | | 0.006 | 0.297 | < 0.001 | < 0.001 |
| b-type heme | | | | 0.002 | < 0.001 | < 0.001 |
| Divalent cation | | | | | < 0.001 | < 0.001 |
| FAD | | | | | | < 0.001 |
| 4-Hydroxycinnamic acid | | | | | | |

**Supplementary Table S4.** Statistical summary of a distance-based Redundancy Analysis (db-RDA) with Bray-Curtis dissimilarities between the sensor domain repertoires of soil bacterial communities as response variable and environmental factors as explanatory variables (N = 255 metagenomes).

| Response variable | Parameters | F-value | P-value | Overall model significance |
|---|---|---|---|---|
| Bacterial sensor domain repertoire | Net Primary Productivity | 16.76 | < 0.001 | F = 8.77<br>P < 0.001 |
| | Conductivity | 3.80 | 0.009 | |
| | Organic carbon | 2.44 | 0.046 | |
| | pH | 11.21 | < 0.001 | |

**References**

1. Bissett A et al. Introducing BASE: the Biomes of Australian Soil Environments soil microbial diversity database. *Gigascience* 2016;**5**:s13742-016.

2. Gumerov VM et al. Amino acid sensor conserved from bacteria to humans. *Proc Natl Acad Sci* 2022;**119**:e2110415119.

3. Cerna-Vargas JP et al. Amine-recognizing domain in diverse receptors from bacteria and archaea evolved from the universal amino acid sensor. *Proc Natl Acad Sci* 2023;**120**:e2305837120.

4. Monteagudo-Cascales E et al. Ubiquitous purine sensor modulates diverse signal transduction pathways in bacteria. *Nat Commun* 2024;**15**:5867.

5. Pardoux R, Dolla A, Aubert C. Metal-containing PAS/GAF domains in bacterial sensors. *Coord Chem Rev* 2021;**442**:214000.

6. Henry JT, Crosson S. Ligand-binding PAS domains in a genomic, cellular, and structural context. *Annu Rev Microbiol* 2011;**65**:261–286.
